# ULK1 drives NDP52-mediated selective autophagic degradation of MHC-I to promote immune evasion in HPV-positive head and neck cancer

**DOI:** 10.64898/2026.03.14.711071

**Authors:** Lexi Vu, Nicholas S. Giacobbi, Mohamed I. Khalil, Canchai Yang, William J. Eckerman, Evelyn Gomez Recinos, Johnathon D. Garber, Hyomin Son, Prerna Chahal, Dara M. Villa, Tasha Srivastava, Abigail Z. Bennett, Katie R. Martin, Craig C. Welbon, Caitlin Williamson, William C. Spanos, Jeffrey P. MacKeigan, Andrew J. Olive, Dohun Pyeon

## Abstract

Antigen presentation by major histocompatibility complex class I (MHC-I) is critical for tumor cell killing by CD8^+^ T cells. Accordingly, tumor cells downregulate MHC-I expression through multiple mechanisms, thereby evading the immune response. Importantly, lower levels of MHC-I are associated with poor responses to immune checkpoint inhibitor therapy. Our recent study has shown that the human papillomavirus (HPV) oncoproteins induce MHC-I protein ubiquitination by membrane-associated Ring-CH-type finger 8 (MARCHF8) in HPV-positive head and neck cancer (HPV+ HNC). However, the mechanism by which ubiquitinated MHC-I is degraded remains elusive. By performing genome-wide CRISPR screens, we identified components of the ULK1 and PIK3C3 complexes for autophagy initiation complexes among the top negative regulators of cell-surface MHC-I expression in HPV+ HNC cells. We show that MHC-I is recruited from the ER to autophagosomes by the cargo receptor NDP52, decreasing MHC-I levels. Further, inhibiting the initiation or nucleation steps of autophagy before autophagosome formation is critical for restoring MHC-I levels on the cell surface. Finally, genetic inhibition of autophagy initiation suppresses HPV+ HNC tumor growth *in vivo* and enhances the CD8^+^ T cell-mediated antitumor response. Our findings suggest that autophagic degradation of MARCHF8-ubiquitinated MHC-I is a key immune evasion mechanism in HPV+ HNC.

## INTRODUCTION

Human papillomavirus (HPV) is the causative agent of ∼5% of all human cancers, with an annual incidence of ∼900,000 cases worldwide^1,2^. HPV-driven carcinogenesis requires the establishment of persistent infection and evasion of the host immune response, facilitated in part by the virally induced immunosuppressive tumor microenvironment (TME)^3^. One of the most common immune evasion mechanisms is downregulating major histocompatibility complex class I (MHC-I) expression on the tumor cell surface. MHC-I presents epitope peptides, including viral antigens and tumor neoantigens, to CD8^+^ T cells in HPV-positive head and neck cancer (HPV+ HNC)^4,5^. CD8^+^ T cell recognition of antigens presented by MHC-I leads to tumor cell killing. Accordingly, the number of tumor-infiltrating CD8^+^ T cells in the TME correlates with improved survival in patients with head and neck cancer^6–9^.

Chemotherapy, radiation, and surgery are effective in treating HPV+ HNC, but they frequently lead to serious adverse effects that can significantly impair the patient’s quality of life. Immune checkpoint inhibitors (ICIs) are a promising alternative that work to reactivate exhausted T cells. However, despite higher levels of tumor-infiltrating T cells and PD-L1 on tumor cells in HPV+ HNC compared to HPV-negative HNC (HPV− HNC), two clinical trials evaluating anti-PD-1/PD-L1 immunotherapies have shown no significant difference in efficacy between HPV+ and HPV− HNC patients^10,11^. Previous studies have suggested that limited MHC-I antigen presentation on tumor cells is a key factor driving the lack of patient responses to ICIs in various cancers^12–14^. However, the mechanisms underlying MHC-I downregulation in HPV+ HNC cells are largely unknown. Therefore, understanding how MHC-I is downregulated in HPV+ HNC is critical for improving patient responses to ICIs.

We recently found that HPV induces the expression of membrane-associated Ring-CH-type finger 8 (MARCHF8), which ubiquitinates MHC-I for degradation^15^. MARCHF8 is a member of the MARCHF subfamily of RING-CH E3 ubiquitin ligases and was first discovered as the human homolog of Kaposi’s Sarcoma-associated herpesvirus (KSHV) proteins K3 and K5^16^. Ubiquitination by MARCHF8 mediates degradation of immune receptors, including MHC-I and the TNF receptor superfamily death receptors FAS, TRAILR1, and TRAILR2^17–19^. As these immune receptors play an important role in antitumor immune responses, the upregulation of MARCHF8 by the HPV oncoproteins is likely to contribute to immune evasion mechanisms and HPV+ HNC progression.

Autophagy is a cellular catabolic process that eliminates and recycles cellular components. It was initially characterized as a bulk degradation process that degrades cytoplasmic proteins and organelles by forming double-membrane vesicles, known as autophagosomes, in response to nutrient starvation^20–22^. However, subsequent studies have demonstrated that proteins and organelles can be selectively degraded by autophagy, even under nutrient-replete conditions^23–25^. This process, known as selective autophagy, requires various cargo receptors that specifically recognize and redirect ubiquitinated target proteins to autophagosomes. These selective autophagy receptors (SARs) contain a ubiquitin-associated (UBA) domain and an LC3-interacting region (LIR)^23,26^. Upon recognition of target proteins by SARs, the Unc51-like kinase (ULK1) complex initiates autophagosome formation and induces autophagy. Next, autophagosomes form via the lipidation of light chain 3B (LC3B), a ubiquitination-like process involving autophagy-related proteins (ATGs)^27–29^. Finally, the mature autophagosome fuses with lysosomes, where its cargo is degraded by lysosomal hydrolases within the acidified lumen. Several previous studies have suggested that autophagy contributes to cancer immune evasion and that inhibition of autophagy can enhance patient responses to immunotherapies^30–33^.

Here, we report that MARCHF8-ubiquitinated MHC-I is recognized by the SAR NDP52 (Nuclear Dot Protein 52 kDa) and degraded through selective autophagy in HPV+ HNC cells. Further, inhibition of autophagy initiation, but not the later steps of autophagosome formation and lysosomal degradation, restores MHC-I expression on the cell surface, activates CD8^+^ T cells, and inhibits tumor growth *in vivo*. These findings provide a novel mechanism of virus-mediated cancer immune evasion in HPV+ HNC, whereby the virus degrades MHC-I proteins via selective autophagy.

## RESULTS

### Genome-wide CRISPR screens identify autophagy genes as top negative regulators of MHC-I expression on HPV+ HNC cells

We have previously shown that HLA-A/B/C protein levels are decreased in HPV+ cell lines compared to normal keratinocytes^15^. To determine HLA-A/B/C levels in HPV+ HNC patients, we first examined HLA-A, HLA-B, and HLA-C mRNA levels in our previously published gene expression dataset (GSE6791)^34^. Surprisingly, our results indicated that mRNA expression of all three alpha chains was significantly upregulated in HPV+ HNC patients compared to normal tissues (Fig. 1A-1C). These findings align with a prior observation from the Cancer Genome Atlas (TCGA) dataset^35^. Next, HLA-A/B/C protein levels were determined by immunofluorescence microscopy using 23 HPV+ HNC, 11 HPV-negative (HPV-) HNC, and 16 normal tonsillar tissue samples (Table S1). Interestingly, HLA-A/B/C fluorescence intensity was significantly lower in both HPV+ and HPV− HNC tumor cells than in normal epithelial cells, as identified by pan-cytokeratin staining and confirmed by hematoxylin and eosin staining (Fig. 1D and S1A). Our results suggest that MHC-I expression in HPV+ and HPV− HNC is downregulated at the protein level.

**Fig. 1.**
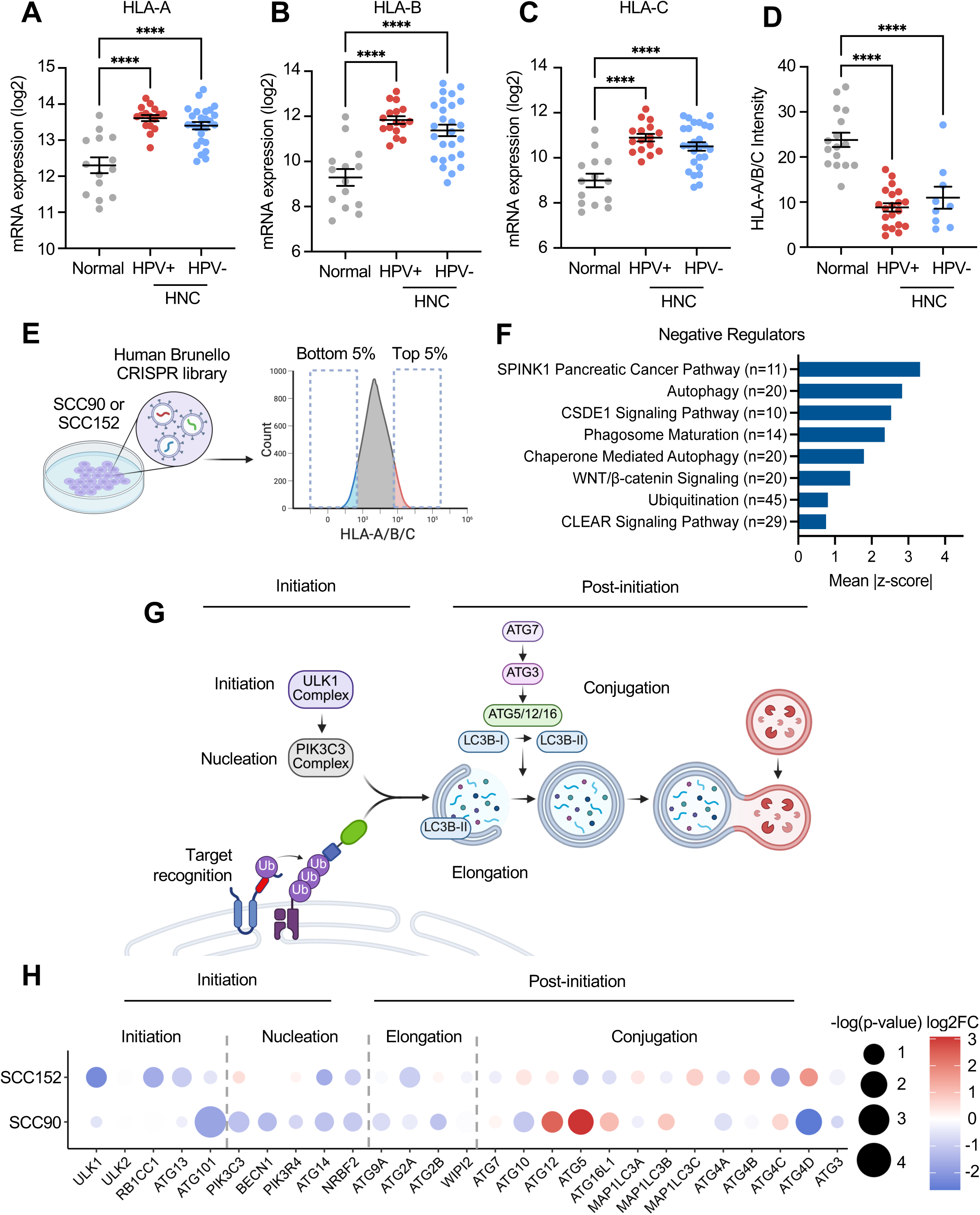
Genome-wide CRISPR screens in HPV+ HNC cells identify autophagy initiation genes as negative regulators of MHC-I expression. mRNA levels of HLA-A (A), HLA-B (B), and HLA-C (C) from microdissected human tonsillar and oral cavity tissue samples from normal individuals (n=12), HPV+ HNC (n=16), and HPV− HNC (n=26) patients (GSE6791). Tissue sections of normal tonsil (n=16), HPV+ HNCs (n=23), and HPV− HNCs (n=11) were stained with hematoxylin & eosin, anti-HLA-A/B/C, anti-pan-cytokeratin, and DAPI. HLA-A/B/C intensity was measured in cytokeratin-positive areas using NIH ImageJ (D). *P*-value was determined by one-way ANOVA analysis, and the mean & standard error of the mean are shown. ****p<0.0001. SCC90 and SCC152 cells expressing Cas9 were transduced with the human Brunello CRISPR library. Cells were sorted for the top 5% and bottom 5% of MHC-I expression levels (E). SgRNA abundance was determined by deep sequencing, and ranked lists of negative regulators of MHC-I expression were identified using the Model-based Analysis of Genome-wide CRISPR-Cas9 Knockout (MAGeCK) software (F). A schematic of the autophagy pathway highlights the target recognition, initiation, nucleation, elongation, and conjugation steps (G). The dot plot shows genes involved in autophagy initiation or post-initiation, as identified by CRISPR screens in SCC90 and SCC152 cells. Dot size represents −log10(p-value) and color represents log2 fold change, with red indicating a stronger positive regulator and blue representing a stronger negative regulator (H).

To identify key factors regulating MHC-I levels on HPV+ HNC cells, we performed genome-wide CRISPR knockout screens using the Brunello small guide RNA (sgRNA) library in two HPV+ HNC cell lines (SCC90 and SCC152) expressing Cas9, as previously described^36^. HPV+ HNC cells with the top and bottom 5% MHC-I levels on the cell surface were isolated by cell sorting. Deep sequencing was then used to quantify sgRNA abundance in the cells of each sorted population (Fig. 1E, Table S2). We then ranked genes using the Model-based Analysis of Genome-wide CRISPR/Cas9 Knockout (MAGeCK) algorithm, which is based on several factors, including the reduction or enrichment of sgRNAs in the lowest 5% MHC-I (MHC-I-low) and the highest 5% MHC-I (MHC-I-high) expressing cells, in addition to the agreement between independent sgRNAs targeting the same gene^37^. Candidate genes that were significantly enriched in MHC-I-low and MHC-I-high groups (a >2-fold change and a p-value <0.05) were considered as positive and negative regulators of MHC-I expression, respectively.

To identify the pathways that positively and negatively regulate MHC-I, we performed a core analysis in Ingenuity Pathway Analysis using the ranked gene list. In both the SCC90 and SCC152 screens, sgRNAs targeting critical components of the antigen presentation pathway were identified as significant positive regulators of MHC-I, confirming the robustness of the screens (Fig. S2A). To further validate the screen results, we generated *TAPBP* knockout cells (SCC90 and SCC152) using two distinct sgRNAs. Knockout efficiency was determined by genome sequencing using Tracking of Indels by Decomposition (TIDE) analysis^38^. A threshold of >30% indel frequency, an R^2^ value >0.9, and a background signal of aberrant sequences <10% were used to define efficient knockout^39–41^. We then determined HLA-A/B/C protein levels using flow cytometry and found that TAPBP knockout significantly decreased HLA-A/B/C expression (Fig. S2B and S2C), confirming the validity of the screens.

Interestingly, the top pathways among negative regulators include autophagy, autophagosome formation, phagosome maturation, and ubiquitination (Fig. 1F, S1D, and S1E, Table S3), all of which are related to protein degradation and autophagy. The autophagic degradation of proteins and cargo can be further categorized into autophagy initiation and post-initiation (Fig. 1G). Autophagy is initiated by the ULK1 complex and membrane nucleation by the PIK3C3 complex, followed by post-initiation steps that encompass autophagosome elongation, conjugation, and autophagosome-lysosome fusion. Upon further dissecting the autophagy genes responsible for these steps, the genes involved in autophagy initiation and nucleation showed the strongest phenotypes as compared to the post-initiation genes (Fig. 1H). These results suggest that autophagy initiation is the key step in preventing MHC-I cell-surface expression in HPV+ HNC cells.

### Inhibiting autophagy initiation is critical for restoring cell-surface MHC-I levels in HPV+ HNC cells

To validate the genes from the CRISPR screen and determine if autophagy initiation factors are negative regulators of MHC-I, we established stable SCC90 and SCC152 cell lines with knockout of the genes in the ULK complex (*ULK1, ULK2, RB1CC1*, *ATG13, ATG101*) and PIK3C3/VPS34 (*BECN1, PIK3R4, ATG14*, *NRBF2*) complex^42^ using two unique sgRNAs. Flow cytometry and western blot analyses showed that knocking out components of the ULK1 and PIK3C3 complexes significantly increased MHC-I levels on HPV+ HNC cells (Fig. 2A – 2F, S3A – S3E). Next, we knocked out the genes required for autophagosome formation (*ATG5* and *ATG7*) after autophagy initiation in SCC90 and SCC152 cells. Knockout efficiency was validated by measuring protein levels of ATG5 and ATG7, along with LC3B as an autophagosome marker^43^ (Fig. 2G, S3F). Either *ATG5* or *ATG7* knockouts showed an increase in total HLA-A/B/C protein levels (Fig. 2G and S3F), but not in cell-surface HLA-A/B/C protein levels (Fig. 2H, 2I, S3G, and S3H), further supporting that inhibiting autophagy post-initiation does not restore cell-surface MHC-I levels on HPV+ HNC cells.

**Fig. 2.**
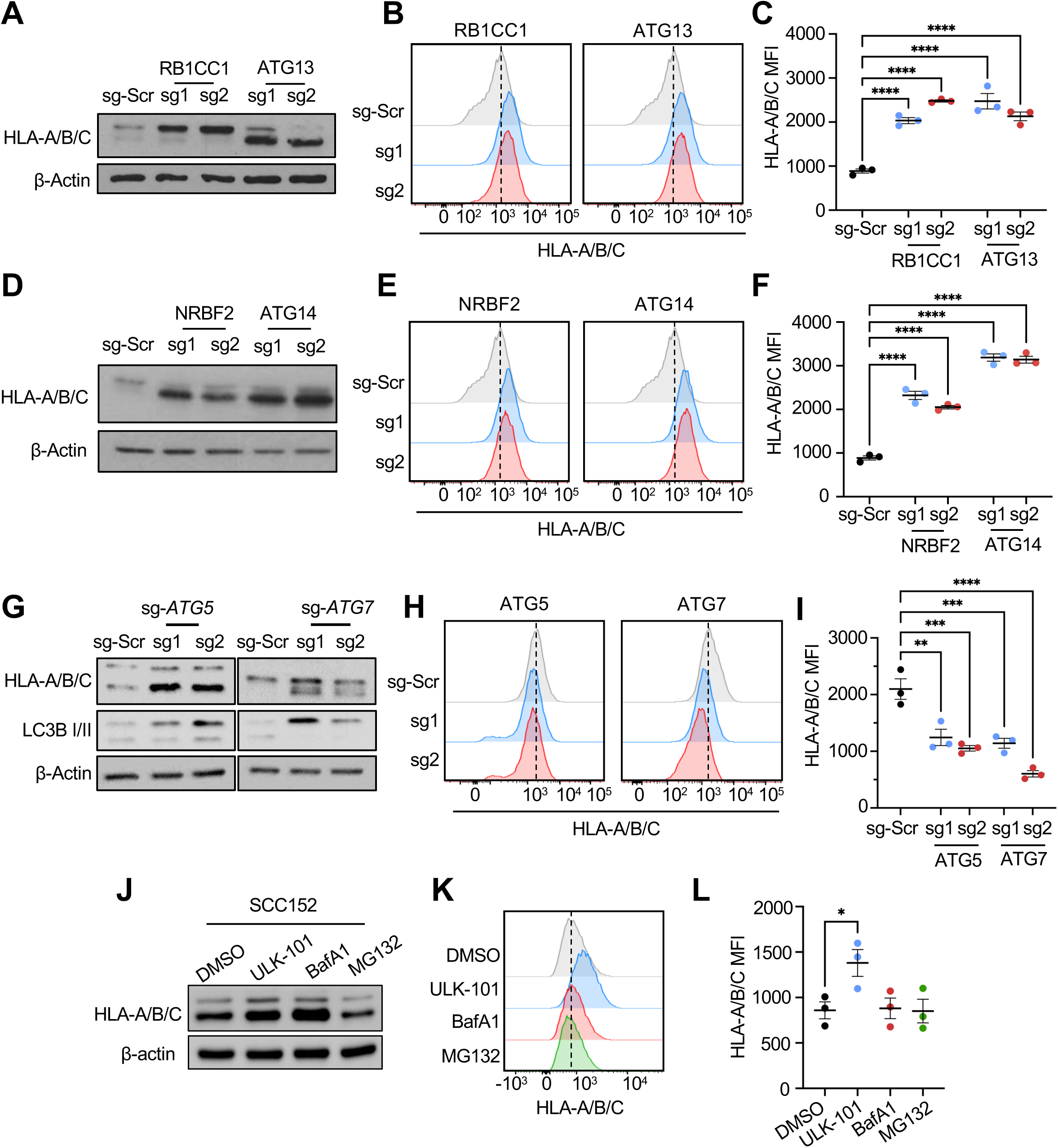
Genetic and pharmacological inhibition of autophagy initiation, but not post-initiation, restores surface MHC-I expression. Genes related to the ULK1 and PIK3C3 complexes were validated via individual CRISPR/Cas9 knockout of indicated genes using two independent sgRNAs in SCC152 cells. Total and cell surface HLA-A/B/C expression was evaluated by western blotting (A and D) and flow cytometry (B, C, E, and F), respectively. Genes involved in autophagosome formation, ATG5 and ATG7, were knocked out using two independent sgRNAs in SCC152 cells. Total and cell surface HLA-A/B/C expression was evaluated by western blotting (G) and flow cytometry (H and I), respectively. SCC152 cells were treated with inhibitors against autophagy initiation (5 µM ULK-101), post-initiation (100 nM Bafilomycin A1), and the proteasome (10 µM MG132) for 16 h. Total and cell surface MHC-I were determined by western blotting (J) and flow cytometry (K and L), respectively. *P*-value was determined by One-way ANOVA. \**p* < 0.05, \*\**p* < 0.01, \*\*\**p* < 0.001, ****p < 0.0001.

We next assessed whether pharmacological inhibition of autophagy initiation restores surface MHC-I expression. We treated SCC90 and SCC152 cells with the potent and selective ULK1 inhibitor ULK-101^44^, the vacuolar H+-ATPase inhibitor Bafilomycin A1 (BafA1) that prevents lysosomal degradation, and the proteasome inhibitor MG132, and measured total and cell-surface HLA-A/B/C protein levels by western blot and flow cytometry, respectively. ULK-101 has been previously tested in cancer cell models; we sought to validate whether treatment of HPV+ HNC cells with ULK-101 inhibited autophagosome formation. SCC152 cells were transduced with the classic autophagosome marker, microtubule-associated protein 1 light chain 3 beta (MAP1LC3B) fused to GFP (LC3B-GFP) to visualize autophagosomes. Cells were then treated with either vehicle (DMSO), BafA1 (100 nM), or BafA1 and ULK-101 (5 µM), and LC3B puncta were visualized by immunofluorescence (Fig. S4A). As expected, treatment with BafA1 resulted in significant accumulation of LC3B-positive puncta relative to controls, an effect that was abolished by addition of ULK-101 (Fig. S4B). This confirms that the ULK1/2 serine threonine kinase inhibitor ULK-101 inhibits autophagy in HPV+ HNC cells. Interestingly, treating SCC90 and SCC152 cells with MG132 did not increase total MHC-I protein levels (Fig. 2J – 2L, S3I – S3K). In contrast, both BafA1 and ULK-101 increased total MHC-I protein while only the ULK-101 treatment increased cell surface MHC-I protein levels (Fig. 2K, 2L, S3J, and S3K). We next treated SCC152 cells with ULK-101, chloroquine (CQ), another post-initiation autophagy inhibitor, or BafA1, and assessed MHC-I expression. Our results show a significant time-dependent increase in total and surface MHC-I levels in ULK-101-treated SCC152 cells (Fig. S4C, S4D), but not BafA1- or CQ-treated cells (Fig. S4E, S4F, S4G, S4H). Taken together, these results suggest that inhibiting autophagy before autophagosome formation is critical for restoring cell-surface MHC-I levels on HPV+ HNC cells.

### The selective autophagy receptor NDP52 binds to and recruits MHC-I protein to autophagosomes

A subset of ubiquitinated proteins is recognized and recruited to the autophagosomes by specific SARs for autophagic degradation. Interestingly, our genome-wide CRISPR screens identified neighbor of BRCA1 (NBR1), sequestosome 1 (*SQSTM1*/p62), and nuclear dot protein 52 (*CALCOCO2*/NDP52) as potential negative regulators of MHC-I expression on HPV+ HNC cells. To validate whether these SARs contribute to the recruitment of MHC-I to autophagosomes for degradation, *CALCOCO2*/NDP52, *NBR1*, and *SQSTM1*/p62 were knocked out in HPV+ HNC cells (SCC152) using two unique sgRNAs each (Tables S4, S5, and S6). Knockouts of NDP52 or NBR1 significantly increased MHC-I protein levels on the cell surface, while p62 knockout did not change MHC-I protein levels (Fig. 3A, 3B, and 3C). Next, to determine which SARs directly bind to MHC-I protein, we pulled down MHC-I proteins (HLA-A/B/C) in SCC152 cells treated with BafA1 and detected NDP52, NBR1, and p62 proteins by western blotting. The results showed that NDP52, but not NBR1 and p62, was pulled down with MHC-I proteins (Fig. 3D), suggesting that NDP52 interacts with MHC-I. Further, we show that co-localization of MHC-I with NDP52 is significantly higher than with p62 using confocal microscopy (Fig. 3E and 3F). Taken together, these results suggest that NDP52 plays an important role in recruiting MHC-I proteins to autophagosomes in HPV+ HNC cells.

**Fig. 3.**
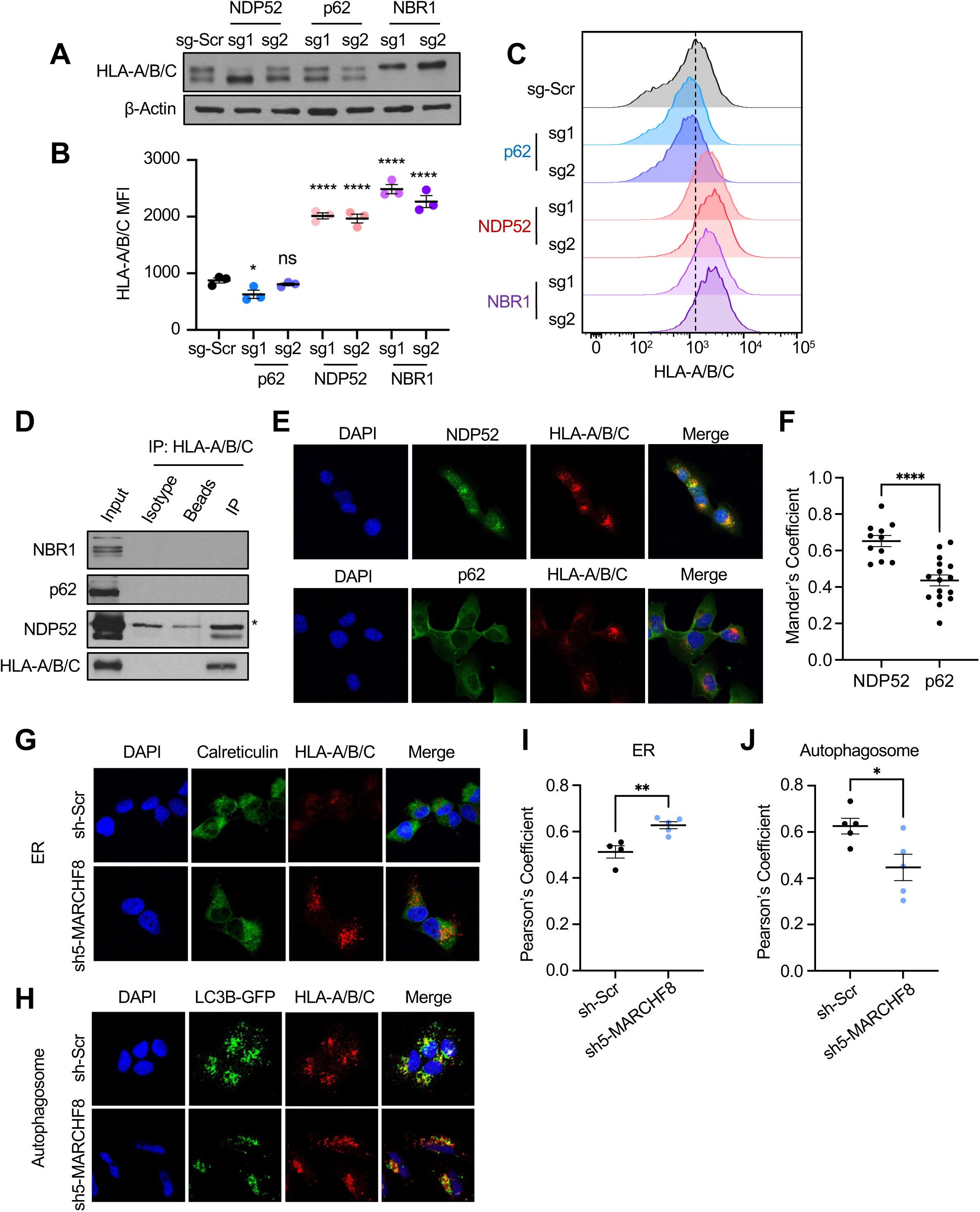
MHC-I is recognized for autophagosome degradation by the selective autophagy receptor NDP52. Autophagy receptor genes (*CALCOCO2*/NDP52, *SQSTM1*/p62, NBR1) were knocked out using CRISPR/Cas9 in SCC152 cells. Total (A) and cell surface (B and C) HLA-A/B/C were determined by western blotting and flow cytometry, respectively. HLA-A/B/C protein was pulled down from lysates of SCC152 cells treated with BafA1 using an anti-HLA-A/B/C antibody, and the autophagy receptors were detected via western blotting. Asterisk (*) denotes nonspecific band. Co-localization of MHC-I with NDP52 and p62 was visualized using immunofluorescence (E). Mander’s coefficient to quantify the overlap of MHC-I with either NDP52 or p62 was calculated in NIH ImageJ (F). Co-localization of MHC-I with calreticulin (ER) (G) or LC3B-GFP (autophagosome) (H) was evaluated in SCC152 cells with sh-Scr or sh5-*MARCHF8* via immunofluorescence. Co-localization was determined by quantifying overlapping colors using NIH ImageJ (I and J). All data shown are mean ± SEM and representative of at least 3 independent experiments. *P*-value was determined using Student’s *t-*Test. *p < 0.05, **p < 0.01, ***p < 0.001, ****p < 0.0001.

The SARs recognize target proteins by detecting specific ubiquitin linkages. Since we previously found that MARCHF8 ubiquitinates MHC-I protein for degradation^15^, we tested whether MARCHF8-mediated ubiquitination recruits MHC-I protein from the ER to autophagosomes. We determined if MARCHF8 knockdown alters the subcellular localization of MHC-I proteins in the ER (calreticulin as a marker) versus autophagosomes (LC3B-GFP as a marker) in HPV+ HNC cells using confocal microscopy. MARCHF8 knockdown increased co-localization of MHC-I with calreticulin (Fig. 3G and 3I) but decreased with LC3B-GFP (Fig. 3H and 3J). These results suggest that MARCHF8-mediated ubiquitination directs MHC-I proteins from the ER to autophagosomes, likely through NDP52 interactions.

### *Ulk1* knockout suppresses HPV+ HNC tumor growth in vivo

Based on our findings, we hypothesize that blocking autophagy initiation enhances MHC-I antigen presentation and antitumor immunity in vivo. To determine if deleting *Ulk1* in HPV+ HNC cells suppresses tumor growth in vivo, we generated two *Ulk1* knockout HPV+ HNC cell lines with the mouse oral epithelial cell line mEERL expressing the HPV oncoprotein E6 and E7 and mutant H-Ras (mEERL/sg2-*Ulk1* and mEERL/sg3-*Ulk1*). *Ulk1* knockout was validated by western blotting for total and phospho-ATG14, a direct ULK1 phosphorylation substrate. Both *Ulk1* knockouts showed a complete absence of phospho-ATG14, with no significant difference in total ATG14 protein levels (Fig. 4A). Cell surface expression of MHC-I was evaluated by flow cytometry using antibodies against H-2Db and H-2Kb, two mouse MHC-Iα haplotypes in C57BL/6J mice. Both H-2Db and H-2Kb showed significant upregulation on the cell surface following *Ulk1* knockout (Fig. 4B and 4C). To determine the effect of *Ulk1* knockout on tumor growth, mEERL/sg-Scr or mEERL/sg-*Ulk1* cells were injected into C57BL/6J mice (n=9/group). All mice injected with mEERL/sg-Scr cells rapidly grew tumors and succumbed to tumor burden within seven weeks post tumor cell injection. Strikingly, both sg2-*Ulk1* and sg3-*Ulk1* syngeneic tumors failed to grow within the same period (Fig. 4D). In fact, two out of nine mice injected with mEERL/sg2-*Ulk1* cells remained tumor-free up to 10 weeks post-injection, and seven mice showed regressed tumors, three of which showed complete tumor regression. Only one mouse injected with mEERL/sg3-*Ulk1* cells formed a palpable tumor, while the rest remained tumor-free for up to 10 weeks. Overall survival post-tumor challenge was monitored, and significant differences were observed in the ULK1 loss groups (Fig. 4E). These results suggest that ULK1-mediated autophagy initiation is necessary for HPV+ HNC tumor growth.

**Fig. 4.**
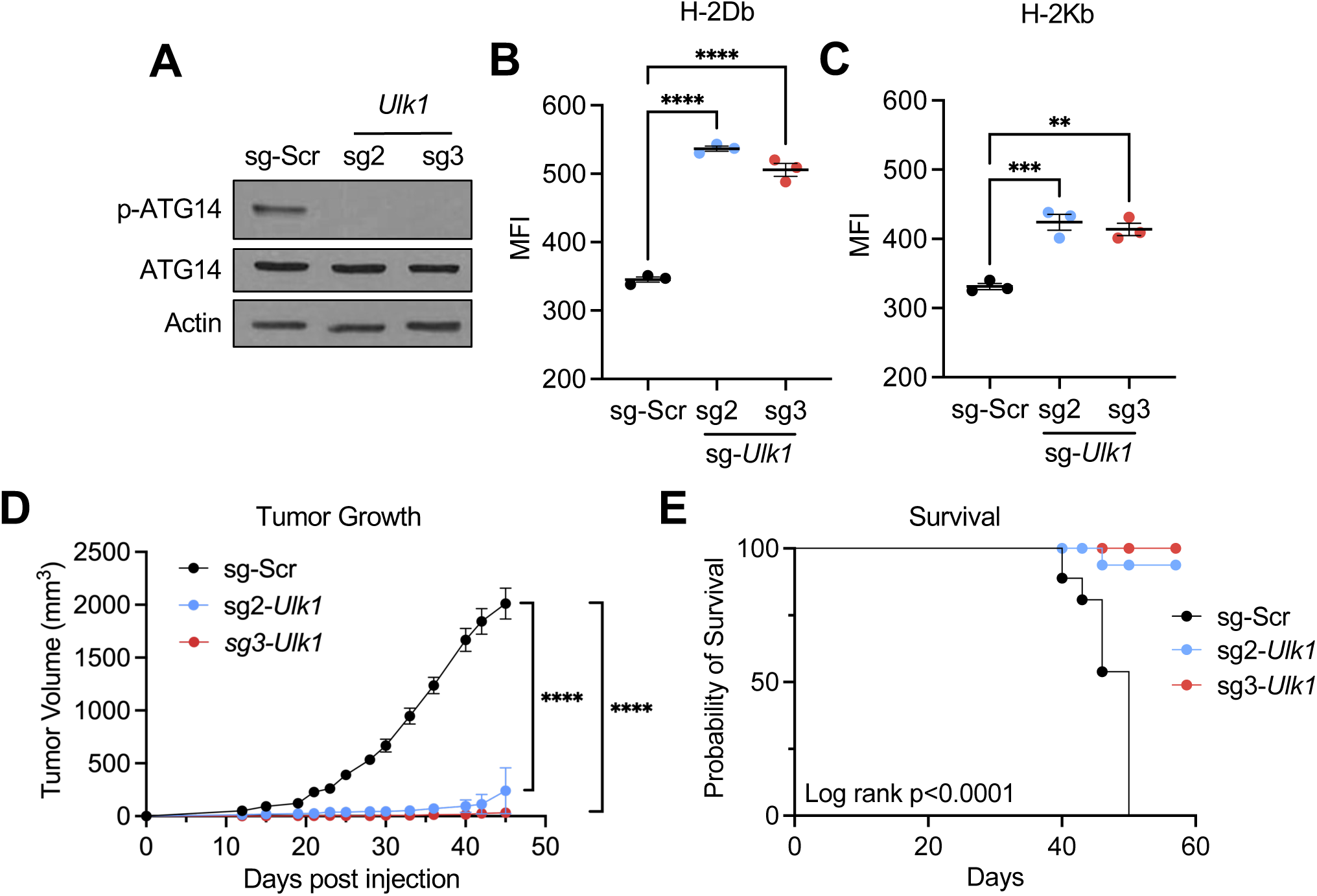
Knockout of *Ulk1* significantly increases MHC-I expression in mouse HPV+ HNC cells and suppresses tumor growth in vivo. *Ulk1* was knocked out from mouse oral epithelial cells expressing HPV16 E6, HPV16E7, and mutant H-Ras (mEERL) using CRISPR/Cas9 and two *Ulk1* sgRNAs (sg2-*Ulk1* and sg3-*Ulk1*). Knockout was validated by western blotting with antibodies that detect total and phospho-ATG14. β-actin was used as a loading control (A). Cell-surface MHC-I expression was assessed by flow cytometry for the two mouse MHC-I haplotypes, H-2Db and H-2Kb (B and C). 5×10^5^ mEERL cells were injected into the right flank of C57BL/6 mice (n=9/group). Tumor growth was measured at least twice per week (D). Mouse survival was estimated using a Kaplan-Meier estimator (E). *P*-value was determined by One-way ANOVA. ****p < 0.0001.

### *Ulk1* knockout enhances CD8^+^ T cell-mediated tumor cell killing

To determine the mechanism of tumor suppression by *Ulk1* knockout, we performed immune cell profiling on tumor tissues from C57BL/6 mice at 21 days after injection with either mEERL/sg-Scr or mEERL/sg2-*Ulk1* cells (Table S7, Fig. S5). Because no tumors developed in mice injected with mEERL/sg3-*Ulk1* cells, we were unable to analyze immune cells. Our results showed that the percentage of CD45^+^ cells decreased in the mEERL/sg2-*Ulk1* tumors compared to mEERL/sg-Scr tumors (Fig. S6A). Of the immune cell subsets assayed (Fig. S6B), total T cells (Fig. 5A), CD8^+^ T cells (Fig. 5B), and macrophages (Fig. S6C) were significantly increased. NK cell (Fig. S6D), B cell (Fig. S6E), and type 1 dendritic cells (cDC1s) (Fig. S6G) showed no significant difference between the two groups, whereas type 2 dendritic cells (cDC2s) (Fig. S6F) were significantly decreased. Interestingly, subtypes of myeloid-derived suppressor cells (MDSCs) show opposite effects: monocytic MDSCs increased (Fig. S6H), whereas granulocytic MDSCs decreased (Fig. S6I). Nine out of ten mice injected with mEERL/sg3-*Ulk1* cells remained tumor-free; therefore, tumor-draining lymph nodes (TDLNs) were collected for T cell profiling. Interestingly, the percentages of CD3^+^/CD45^+^ cells significantly decreased between TDLN from mEERL/sg-Scr and mEERL/sg-*Ulk1* mice (Fig. 5D). However, mice injected with mEERL/sg-Scr cells showed a significant decrease in CD8^+^ T cells and a significant increase in CD4^+^ T cells in the TDLNs compared to mice injected with mEERL/sg-*Ulk1* (Fig. 5E and 5F). Within the CD8^+^ T cell population, cells were further stained with anti-CD44 and CD62L antibodies to identify naïve and effector memory T cells. Our results showed that compared to mice injected with mEERL/sg-Scr cells, mice injected with mEERL/sg-*Ulk1* cells had increased effector T cell populations and decreased naïve T cell populations (Fig. 5G and 5H). These results suggest that *Ulk1* knockout enhances T cell and macrophage infiltration into the TME and increases effector and central memory T cells in the TDLNs, thereby enhancing the antitumor CD8^+^ T cell response.

**Fig. 5.**
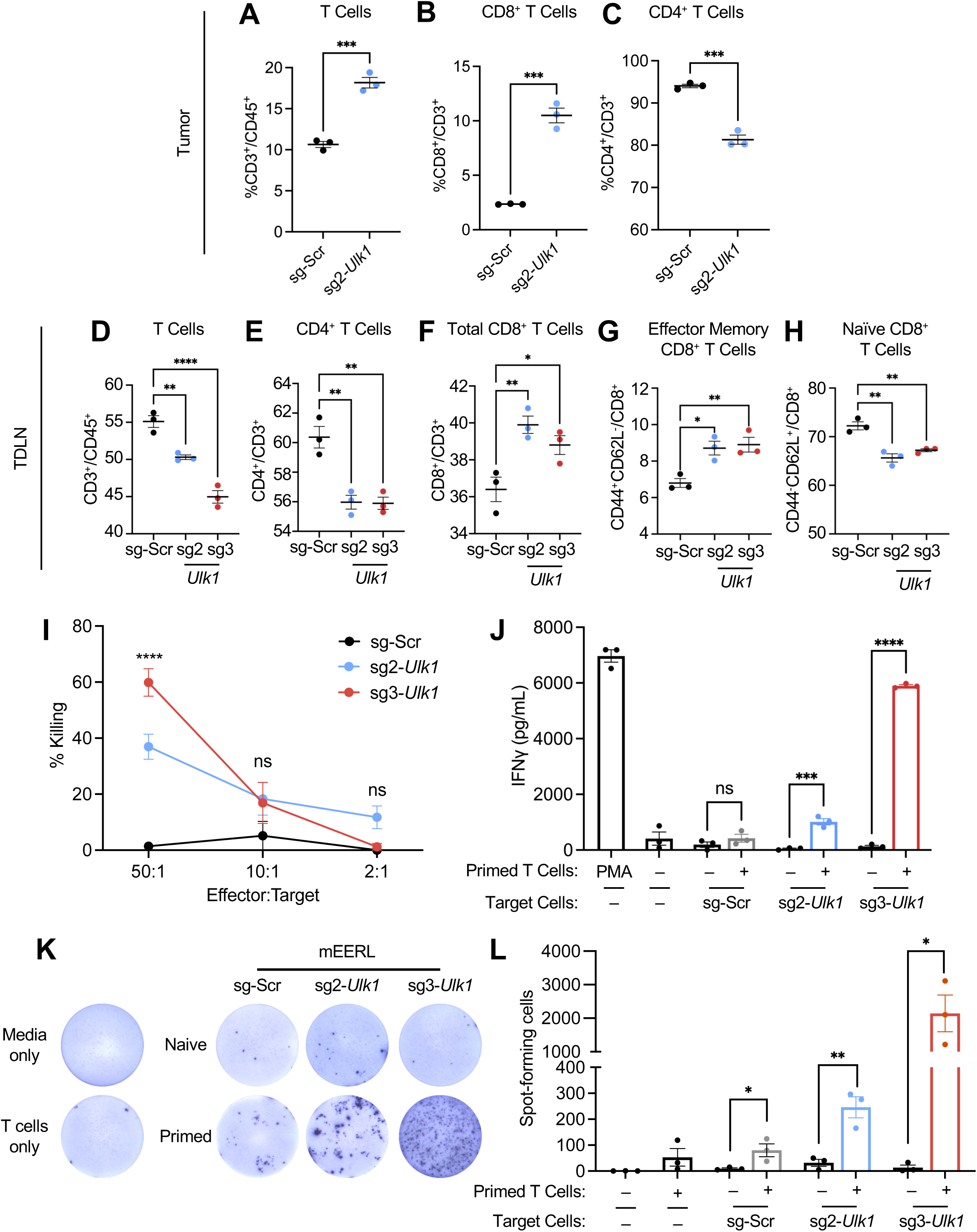
*Ulk1* knockout increases CD8^+^ tumor cell infiltration and enhances CD8^+^ effector functions. Immune cell profiling of tumor tissues from C57BL/6 mice injected with mEERL/sg-Scr or mEERL/sg2-*Ulk1* (n=4/group). Isolated single cell suspensions were stained with an antibody cocktail and analyzed by flow cytometry. Percentages of total T cells of CD45^+^ (A), CD8^+^ T (B), and CD4^+^ T cells (C) of total CD3^+^ cells are shown. T cell profiling of tumor draining lymph nodes (TDLNs) from mice injected with either mEERL/sg-Scr or mEERL/sg-*Ulk1* was isolated 21 days post-injection (n=3/group). Single-cell suspensions were stained with an antibody cocktail containing Zombie NIR, anti-CD45, anti-CD3, anti-CD4, anti-CD8, anti-CD44, and anti-CD62L antibodies. Representative bar graphs showing percentages of total T cells (D), CD4^+^ T (E), and CD8^+^ T cells (F), effector memory CD8^+^ T cells (G), and naïve CD8^+^ T cells (H). *P*-value was determined by One-way ANOVA. \**p*<0.05, \*\**p*<0.01, \*\*\**p*<0.001, \*\*\*\**p*<0.0001. Cytolytic activity of isolated CD8^+^ T cells co-cultured with mEERL cells at a 50:1, 10:1, or 2:1 T cell: tumor cell ratios were measured by LDH assay (I). Effector functions of isolated CD8^+^ T cells were evaluated via IFNγ production measured by ELISA (J) and ELISpot (K and L). The number of spot-forming cells was counted using NIH ImageJ. All data shown are mean ± SEM and representative of at least 3 biological replicates. *P*-value was determined using Student’s *t-*Test. *p < 0.05, **p < 0.01, ***p < 0.001, ****p < 0.0001.

Next, to assess whether *Ulk1* knockout in mEERL cells enhances CD8^+^ T cell effector functions, we isolated CD8^+^ T cells from mice, analyzed tumor cell killing using the lactate dehydrogenase release assay, and measured interferon-γ (IFN-γ) production by ELISA and ELISpot. CD8^+^ T cells from mice injected with *Ulk1* knockout cells showed significantly increased tumor cell killing, whereas CD8^+^ T cells from mice injected with sg-Scr cells showed minimal or no tumor cell killing (Fig. 5I). We next assessed activation of the CD8^+^ T cells by measuring IFN-γ production. While we observed no significant difference in IFN-γ production between naïve and primed CD8^+^ T cells co-cultured with mEERL/sg-Scr cells, primed CD8^+^ T cells showed a significant increase in IFN-γ production when co-cultured with mEERL/sg-*Ulk1* cells (Fig. 5J). Similarly, IFN-γ production was significantly increased in co-cultures of primed CD8^+^ T cells with mEERL/sg-*Ulk1* over naïve CD8^+^ T cells when measured by ELISpot (Fig. 5K and 5L). Taken together, these results suggest that blocking autophagy initiation by ULK1 knockout enhances CD8^+^ T cell activity against HPV+ HNC cells.

## DISCUSSION

Limited antigen presentation by tumor cells has been suggested as a key factor underlying the lack of patient responses to ICI therapy^45–47^. This may explain why HPV+ HNC patients often fail to achieve significant therapeutic benefits from ICI therapy despite high PD-L1 expression and increased tumor-infiltrating T cells compared to HPV− HNC patients^10,48–50^. Interestingly, despite increased mRNA levels (Fig. 1A – 1C), MHC-I protein levels decrease significantly in HPV+ HNC patient tumors. Consistent with the results, our recent findings show that MHC-I protein degradation is mediated by ubiquitination via HPV-induced MARCHF8^15,17^. However, the downstream mechanism by which MHC-I is degraded following MARCHF8-mediated ubiquitination remained unknown. Previous studies have shown that MHC-I is degraded via the autophagosome-lysosome pathway across several cancers, including pancreatic cancer and acute myeloid leukemia^33,51–53^. Further, inhibiting autophagy synergizes with ICI therapy and enhances CD8^+^ T cell-mediated tumor cell killing^54^. Consistent with these results, our genome-wide CRISPR screens in HPV+ HNC cells have identified numerous autophagy genes as the top negative regulators of MHC-I expression. Here, we show that in HPV+ HNC, MARCHF8-ubiquitinated MHC-I is recognized by NDP52 and degraded through autophagy. Furthermore, we show that inhibiting autophagy initiation, but not post-initiation, is critical for restoring cell-surface MHC-I levels. Inhibiting the essential autophagy initiator ULK1 activates the CD8^+^ T cell antitumor response, significantly suppressing HPV+ HNC tumor growth in vivo.

Compared with bulk autophagy, which nonspecifically engulfs cytoplasmic components to recycle building blocks and maintain metabolic homeostasis, selective autophagy requires the specific recognition of ubiquitinated proteins by SAR^22^. The widely studied mammalian SARs include NBR1, p62, NDP52, optineurin (OPTN), and Tax1 binding protein 1 (TAXBP1), all soluble proteins that contain a ubiquitin-binding domain (UBA) and an LC3-interacting region (LIR) to tether targets to autophagosomes^28,55–57^. The SARs that target MHC-I appear to be cancer-type-dependent. Yamamoto et al. have demonstrated an NBR1-dependent mechanism in pancreatic ductal adenocarcinoma (PDAC)^33^, whereas Herhaus et al. have identified a novel SAR, IRGQ, in acute myeloid leukemia^51^, both of which result in the autophagic degradation of MHC-I proteins. Here, we report that MHC-I is recruited to autophagosomes specifically by NDP52. While it has been shown that each SAR recognizes specific targets^58–60^, it remains largely unknown how each SAR identifies target proteins for transport to autophagosomes. One critical factor is the diverse ubiquitin linkages. Ubiquitin contains seven lysine residues (K6, K11, K27, K29, K33, K48, and K63), each of which can be used to form distinct ubiquitin chains. The type of linkage between ubiquitin molecules is a key determinant of the downstream fates of ubiquitinated proteins^61^. While K48 is the canonical signal for proteasomal degradation^62^, the other chains are less studied. Interestingly, MARCHF8 has been shown to add both K27- and K29-linked ubiquitin chains to target proteins^63,64^. Further, K27-linked ubiquitin chains added by MARCHF8 can be recognized by NDP52^64^. Thus, further investigation into the types of ubiquitin linkages on MHC-I can provide insight into how MHC-I is recognized by SARs and targeted for degradation across a multitude of cancer types.

Previous studies have shown that *ATG7* knockout and chloroquine treatment restore cell-surface levels of MHC-I in cancers such as PDAC and during viral infections, including influenza and lymphocytic choriomeningitis viruses^33,65,66^, by inhibiting autophagosome formation and autophagosome-lysosome fusion, respectively. In contrast, our results showed that inhibiting autophagosome formation and autophagosome-lysosome fusion fails to restore cell-surface MHC-I levels. Instead, blocking autophagy initiation is critical for restoring cell-surface MHC-I levels on HPV+ HNC cells (Fig. 2). Inhibiting early steps, such as MHC-I targeting or activation of the ULK1 complex, may prevent sequestration of MHC-I molecules into autophagosomes, whereas blocking downstream steps fails to alter the fate of MHC-I once sequestration has occurred. We and others have shown that MARCHF8 predominantly co-localizes in the ER^15,67^, and MARCHF8 knockdown increases MHC-I localization in the ER (Fig. 3F and 3H). Consistently, the ER also serves as a common site for ULK1-mediated autophagy initiation^68,69^. Inhibition of either MARCHF8 or ULK1 restores MHC-I on the cell surface, suggesting that ubiquitinated MHC-I in the ER likely enters autophagosomes rather than being transported to the cell surface. Therefore, targeting autophagy by inhibiting ULK1 could be a promising strategy to activate antitumor T cell responses and treat HPV+ HNC patients.

*Ulk1* knockout in our HPV+ HNC cells resulted in significant tumor suppression and enhanced CD8^+^ T cell-mediated killing of tumor cells (Fig. 4E and 5J). Furthermore, CD8^+^ T cells primed with *Ulk1* knockout tumor cells were able to kill wildtype HPV+ HNC cells (Fig. S7B). These robust anti-tumor effects occurred independently of ULK2, a serine/threonine kinase previously shown to partially compensate for ULK1^70,71^. This suggests a reliance on ULK1, but not ULK2, for autophagy initiation in the mEERL model, as evidenced by the complete loss of phospho-ATG14 following ULK1 knockout in mEERL cells (Fig. 4A). A similar dependency on ULK1 is shown in the human SCC90 and SCC152 cells. While ULK1 appears to be a significant negative regulator in both genome-wide CRISPR screens, ULK2 shows minimal effect on MHC-I surface expression. Nevertheless, ULK2 may still contribute to tumorigenesis through autophagy-independent functions, such as promoting TNF superfamily signaling^72^ and tumor cell invasion^73^. Thus, ULK-101, a potent and selective ULK1/2 inhibitor, could be a promising therapeutic agent for HPV+ HNC patients, particularly those with recurrent or metastatic disease. ULK-101 has demonstrated superior potency and selectivity compared to other ULK1/2 inhibitors^44,74^. Studied initially for the treatment of pancreatic and lung cancers, ULK-101 effectively suppresses autophagy initiation and increases MHC-I levels in HPV+ HNC cells. Additional studies are necessary to determine the in vivo efficacy of ULK-101 in inhibiting autophagy and suppressing tumors.

While inhibition of autophagy to restore MHC-I levels has been shown to be a promising therapeutic strategy, future studies must explore the broader implications of autophagy inhibition for remodeling the TME. Autophagy plays diverse roles in cancer progression, affecting not only tumor cells but also various components in the TME. Indeed, inhibition of autophagy upregulates proinflammatory chemokines, such as CCL5, CXCL10, and IFN-γ, in the TME, thereby reprogramming cold tumors into hot tumors and improving responses to ICI therapies^75^. In addition, inhibition of Beclin1, a downstream substrate of ULK1, promotes NK cell-mediated tumor killing by upregulating the chemokine CCL5 in melanoma^76^. Therefore, future studies are necessary to identify additional autophagy targets within the TME, elucidate their roles in immune regulation, and determine how modulating these targets may affect cancer progression and treatment efficacy. This comprehensive approach will be essential for developing effective combination therapies that leverage autophagy inhibition to enhance antitumor immunity for treating HPV+ HNC patients.

## MATERIALS AND METHODS

### Cell Culture

HPV+ HNC (SCC90 and SCC152) and 293FT cells were purchased from the American Type Culture Collection and Thermo Fisher, respectively. Cells were cultured in Dulbecco’s Modified Eagle’s medium containing 10% fetal bovine serum (FBS) and Antibiotic-Antimycotic (Gibco, No. 15240062). Transformed mouse oral epithelial (mEERL) cells were obtained from Dr. William Spanos and cultured in E-medium (DMEM and F12 media supplemented with 0.005% hydrocortisone, 0.05% transferrin, 0.05% insulin, 0.0014% triiodothyronine, 0.005% EGF, and 2% FBS), as previously described^77^.

### Plasmid Constructs

The pCDH-CMV-LC3B-GFP plasmid construct was prepared as previously described^78^. shRNAs targeting human MARCHF8 were purchased from Sigma-Aldrich. All lentiviruses were produced using the pCMV-VsVg (Addgene #8454) and pCMV-Δ8.2 (Addgene #12263) packaging constructs (gifts from Jerome Schaack).

### Lentiviral Transduction

All lentiviruses were produced using 293FT cells transfected with the packaging constructs and the respective transfer vector using polyethyleneimine (PEI) in a 3:1 PEI:DNA ratio. 72 h post-transfection, cell culture supernatants were collected and centrifuged at 2000xg for 5 min to eliminate cell debris. HPV+ HNC cells were then incubated with the collected supernatants for 48 h in the presence of 15 μg/ml polybrene and selected with 2 μg/ml puromycin or 8 μg/ml blasticidin until a non-transduced control plate showed 100% cell death.

### CRISPR Screen

The human CRISPR Brunello lentiviral library was a gift from David Root and John Doench (Addgene #73179)^36^. The 76,441 sgRNAs in the library targeting 19,114 genes were packaged with pCMV-VsVg and pCMV-Δ8.2 into lentiviruses using 293FT cells. These lentiviruses were transduced into SCC90 and SCC152 cells at an MOI = 0.3. Following puromycin selection (4 µg/mL), cells were fixed, and fluorescence-activated cell sorting (FACS) on a BD S3e cell sorter was used to isolate the top 5% and bottom 5% of MHC-I expressing cells with at least 10^7^ cells per population (>100 X sgRNA coverage). Genomic DNA was extracted from isolated cells and PCR-amplified using P5 and P7 Illumina compatible primers from IDT as previously described^79^ (Table S2). PCR amplicons were sequenced on an Illumina NextSeq 500. After removing adapter sequences, model-based analysis of genome-wide CRISPR-Cas9 knockout (MAGeCK) was used to map the reads to the Brunello library index. The MAGeCK Gene Summary provides a ranked list of genes based on 4 unique sgRNAs, generating a candidate gene list, while the sgRNA summary shows data for each individual sgRNA (GSE324388).

### Generation of CRISPR-targeted Knockouts

To generate CRISPR/Cas9 knockout cell lines, sgRNAs were designed using the web-based software ChopChop^80^ and/or sequences obtained from the human CRISPR Brunello lentiviral pooled library (Table S4). The sgRNAs were synthesized as annealed nucleotides by Integrated DNA Technologies (IDT, Newark, NJ) with overhangs complementary to the sgOPTI-Amp-Puro lentiviral vector (Addgene #85681, a gift from Eric Lander & David Sabatini) for human cell lines and LentiCRISPR v2-Blast (Addgene #83480, a gift from Mohan Babu) for mouse cell lines following BsmBI-v2 restriction digestion. SgRNA-containing plasmid constructs were then packaged into lentiviruses as previously described^17^ and transduced into Cas9-expressing HPV+ HNC cells in the presence of 15 μg/ml polybrene. Following puromycin selection, genomic DNA was extracted, and sgRNA target regions were amplified using PCR and Sanger sequenced (Azenta). Sequencing data were analyzed by Tracking of Indels by Decomposition (TIDE) software^81^ to assess the size and frequency of indels compared to control cells. TIDE primers used are listed in Table S5.

### Flow Cytometry

Flow cytometry was performed as previously described^17^. Briefly, single-cell suspensions were stained with the Zombie NIR Fixable Viability Kit (Biolegend #423105) according to the manufacturer’s protocol, followed by Fc receptor blocking (Biolegend #101301, Biolegend # 422301) and staining with specific antibodies (Table S7). The cells were fixed using 4% paraformaldehyde and analyzed on an Attune CytPix flow cytometer (Thermo Fisher Scientific). Data were analyzed using FlowJo software (Tree Star).

### Immunofluorescence

Formalin-fixed paraffin-embedded (FFPE) human and mouse tissue samples were deparaffinized, rehydrated, and incubated in pH 6.0 10 mM sodium citrate buffer (Sigma-Aldrich, C9999) at 100°C for 20 min for antigen retrieval. Samples were washed in Tris-buffered saline (pH 7.4) + 0.1% Tween-20 (TBS-T), blocked using TBS-T + 1% bovine serum albumin (BSA) + 5% goat serum for 1h at room temperature (RT), and incubated with primary antibodies (Table S7) diluted in blocking buffer at 4°C for 16 h overnight in a humid environment. Samples were then incubated with fluorophore-conjugated secondary antibodies (1:500 in blocking buffer, Table S7) at RT for 1 h, washed with TBS-T, and counterstained with 4’,6-diamidino-2-phenylindole (DAPI) for 5 min. Samples were mounted with ProLong Gold AntiFade Mountant (Invitrogen, P36930) and imaged on a Leica Stellaris 5 Confocal Laser Scanning Microscope (CLSM) using Leica Application Suite X software. Cultured cells were plated on poly-L-lysine-coated coverslips for 48 h and treated with Bafilomycin A1 (BafA1), ULK-101, or MG132 at 37°C for 16 h. Cells were fixed using paraformaldehyde at RT for 15 min, blocked at RT for 1 h using phosphate-buffered saline (PBS) + 0.05% BSA + 0.003% Triton X-100, and incubated at 4°C overnight in primary antibody diluted in PBS + 0.01% BSA + 0.003% Triton X-100 (Table S7). Then, cells were incubated with fluorophore-tagged secondary antibodies (1:500, Table S7) at RT for 1 h, washed with PBS, and counterstained with DAPI for 5 min. Slides were mounted with ProLong Gold AntiFade Mountant (Invitrogen, P36930) and imaged on a Nikon C2 CLSM using NIS-Elements Software.

### Image Analyses

All images were analyzed using NIH ImageJ on grayscale raw image files. HLA-A/B/C intensity in the tumor cells on FFPE tissue sections was measured in cytokeratin-positive areas. Briefly, images were preprocessed to identify regions of interest (ROIs) containing cytokeratin-positive areas. Next, the mean intensity of the HLA-A/B/C signal was measured within the ROIs. LC3B-GFP puncta in cultured cells were quantified using ImageJ. Images were preprocessed by selecting individual cells as ROIs. A difference-of-Gaussians (DoG) filter was applied to enhance puncta visibility and reduce background noise^82^. Manual thresholding was then used to segment puncta from the background. The number of puncta per cell was counted using particle analysis in NIH ImageJ and validated by blind manual counting. P-value was determined by Student’s t-test (p<0.05). Colocalization analyses were performed using the Just Another Co-localization Plugin (JaCoP) in NIH ImageJ. Images were split into individual acquisition channels, and images of interest were used as input into JaCoP. Thresholds were set for each image to identify signal-positive and signal-negative regions, and Pearson’s and Manders’ colocalization coefficient was determined using the JaCoP in NIH ImageJ.

### Immunoprecipitation and Western Blotting

Protein lysates were prepared in 1X radioimmunoprecipitation assay (RIPA) buffer (Sigma) supplemented with a protease inhibitor cocktail (Roche). Protein concentration was determined using the Pierce BCA Protein Assay Kit according to the manufacturer’s instructions (Thermo Scientific, No. 23225). Western blotting was performed with 10-30 μg of whole cell lysates using the antibodies listed in Table S5.

### Mice and Tumor Growth

Animal studies were all performed in compliance with Michigan State University’s Animal Care and Use Committee (IACUC) under protocol PROTO202400214. mEERL cells (5×10^5^ cells/mouse) were subcutaneously injected into the rear right flank of 6- to 8-week-old C57BL/6J mice (Jackson Labs, #000664) (n=10 per group unless otherwise indicated). Tumor lengths (L) and widths (W) were measured with calipers, and volumes (V) were calculated using V = 0.5 x L x W^2^. Tumor tissue and mouse weights were measured at least twice per week beginning on day 12 post-injection.

### Isolation of CD8^+^ T Cells

Mouse spleens were collected in PBS + 2% FBS + 1 mM ethylenediaminetetraacetic acid (EDTA). Spleens were mechanically disrupted using a 5 mL syringe plunger over a 40 µm cell strainer to generate a single-cell suspension. The cell suspension was centrifuged at 350 × g for 10 min. Splenocytes were resuspended in 1X MojoSort Buffer (BioLegend #480017). CD8^+^ T cells were isolated from 1×10^8^ splenocytes using the MojoSort Mouse CD8 T Cell Isolation Kit (BioLegend #480008) according to the manufacturer’s instructions. Isolated CD8^+^ T cells were cultured for 5 days in RPMI 1640 (Gibco) supplemented with 10% FBS, L-glutamine, β-mercaptoethanol, Dynabeads Mouse T-Activator CD3/CD28 (Invitrogen), and recombinant mouse IL-2 (ThermoFisher) every 36 h. 36 h before co-culturing with mitomycin C-treated mEERL cells, CD3/CD28 beads were magnetically removed and replaced with anti-mouse CD28 antibody (BioLegend #102101).

### Lactate Dehydrogenase Assay

Isolated CD8^+^ T cells (effector) were co-cultured with 5,000 mEERL cells (target) with scrambled or *Ulk1* sgRNAs in RPMI1640 containing recombinant mouse IL-2 and anti-mouse CD28 antibody at 50:1, 10:1, or 1:1 effector-to-target ratios. mEERL cells were treated with 1% Triton-X to determine the maximum LDH release, which served as the positive control for 100% killing. CD8^+^ T cells and mEERL cells were plated independently to measure background LDH levels as a negative control. After 18 h, cell cytotoxicity was determined by measuring lactate dehydrogenase production in cell culture supernatant using the Cytotoxicity Detection Kit (Millipore Sigma, No. 11644793001).

### IFN-**γ** ELISA

Isolated CD8^+^ T cells were co-cultured with 5,000 mEERL cells with scrambled or *Ulk1* sgRNAs in RPMI 1640 supplemented with 10% FBS, L-glutamine, β-mercaptoethanol, anti-mouse CD28 antibody, and recombinant mouse IL-2 and anti-mouse CD28 antibody at 50:1 effector-to-target ratios for 72 h. CD8^+^ T cells cultured with phorbol 12-myristate 12-acetate (PMA) and Ionomycin were used as a positive control to determine maximum IFN-γ production. Naïve CD8^+^ T cells alone were cultured independently to determine baseline IFN-γ production. Cell culture supernatants were collected, and IFN-γ production was quantified by ELISA (Biolegend, #430801) according to the manufacturer’s instructions.

### IFN-**γ** ELISpot

Isolated CD8^+^ T cells were co-cultured with 2,000 mEERL cells with scrambled or *Ulk1* sgRNAs in RPMI 1640 supplemented with 10% FBS, L-glutamine, β-mercaptoethanol, anti-mouse CD28 antibody, and recombinant mouse IL-2 and anti-mouse CD28 antibody at 50:1 effector-to-target ratios for 72h in a multiscreen 96-well plate with PVDF membrane (Millipore Sigma, No. MAIPS4510). CD8^+^ T cells cultured with PMA and Ionomycin were used as a positive control. CD8^+^ T cells only were cultured separately to determine baseline IFN-γ production. IFN-γ production was determined using the Mouse IFN-γ ELISpot Development Module (R&D Systems, No. SEL485), visualized using the ELISpot Blue Color Module (Strep-AP and BCIP-BNBT) (R&D Systems, No. SEL002), and imaged using the ImmunoSpot Analyzer (CTL). CTL files were analyzed by NIH ImageJ. A consistent threshold was used to identify spots versus background, with the media-only wells serving as the negative control. Spots were counted using the Analyze Particles function in NIH ImageJ.

### Statistical Analysis

All statistical analyses were performed using GraphPad Prism.

## Data Availability

Raw FASTQ and processed sequencing data is available for download in the NCBI Gene Expression Omnibus (GSE324388). Raw imaging files will be uploaded to Dryad. All data used to generate the graphs and associated p-values is included in the Source Data file.

## ACKNOWLEDGEMENTS

We thank Sandra O’Reilly (IVIS Spectrum Imaging Core), Melinda Frame (Center for Advanced Microscopy), Amy Porter (Investigative Histopathology Laboratory), Matt Bernard, Daniel Vocelle, and Soo Hyun Ahn (Flow Cytometry Core) for technical assistance, and the members of the Pyeon and Wei Wang laboratories for their valuable comments and suggestions. This work was supported by NIH F31 DE034281 (L.V.), NIH R01 DE033797 (D.P.), NIH R01 DE029524 (D.P.), NIH R01 R01CA297993 (J.P.M.), NIH R01 AI165618 (A.J.O), the Henry Ford + MSU Cancer Integration Grant (D.P.), and the MSU Foundation Strategic Partnership Grant (D.P.).

---

**Fig. S1. MHC-I protein levels are decreased in HPV+ and HPV-negative patient tumors.** Tissue sections of normal tonsil (n=16), HPV+ HNCs (n=23), and HPV− HNCs (n=11) were stained with hematoxylin & eosin, anti-HLA-A/B/C (green), anti-pan-cytokeratin (red), and DAPI (blue). Representative images are shown.

**Fig. S2. Genome-wide CRISPR screens identify key regulators of cell-surface MHC-I expression on HPV+ HNC cells.** Pathways enriched in the significant positive regulators (p-value <0.05 and |Log2FC| >1) from the genome-wide CRISPR screens were identified using Ingenuity Pathway Analysis (IPA) (A). Screens were validated by knocking out tapasin (TAPBP) and evaluating HLA-A/B/C expression by flow cytometry (B and C). Volcano plots showing gene hits from the autophagy, autophagosome formation, phagosome maturation, and ubiquitination pathways in SCC90 (D) and SCC152 (E) cells, as identified from Ingenuity Pathway Analysis.

**Fig. S3. Inhibition of autophagy initiation, but not post-initiation, restores surface MHC-I expression in HPV+ HNC cells.** Genes in the ULK1 (*ATG13* and *RB1CC1*) and PIK3C3 (*ATG14* and *NRBF2*) autophagy initiation complexes that were significant negative regulators in the CRISPR screen were knocked out of SCC90 cells. MHC-I surface expression was determined by flow cytometry (A – E). Genes involved in autophagosome formation (*ATG5* and *ATG7*) were knocked out in SCC90 cells. Inhibition of autophagosome formation was assessed by western blotting with antibodies against LC3B-I/II (F). Total (F) and cell-surface (G) MHC-I levels were determined by western blotting and flow cytometry, respectively. SCC90 cells were then treated with inhibitors against autophagy initiation (5 µM ULK-101), post-initiation (100 nM Bafilomycin A1), and the proteasome (10 µM MG132) for 16 h. Total and cell-surface MHC-I levels were determined by western blotting (H) and flow cytometry (I and J), respectively. *P*-value was determined by One-way ANOVA. \**p* < 0.05, \*\**p* < 0.01, \*\*\**p* < 0.001, ****p < 0.0001.

**Fig. S4. Post-initiation autophagy inhibitors do not increase MHC-I over time.** SCC152 cells were transduced with LC3B fused to a GFP construct (LC3B-GFP). Cells were plated on poly-L-lysine-coated glass coverslips and treated with DMSO, 100 nM BafA1, or 100 nM BafA1 plus 5 µM ULK-101 for 16 h. Cells were fixed with 1% paraformaldehyde and imaged on a Keyence Brightfield Microscope (BZ-X710) (A). Puncta were counted using NIH ImageJ (B). *P*-value was determined using Student’s t-test. ****p < 0.0001. Non-significant results are reported as ns. SCC152 cells were then treated with 5 µM ULK-101, 50 µM chloroquine, and 100 nM BafA1 for 0, 2, 4, 8, or 16 h. Total MHC-I was evaluated by western blot. LC3B I/II and phospho-ATG14 were used as markers of autophagy. β-actin was used as a loading control (C, E, and G). Cell surface MHC-I was evaluated by flow cytometry (D, F, and H). *P*-value was determined by One-way ANOVA. \**p* < 0.05, \*\**p* < 0.01, \*\*\**p* < 0.001, ****p < 0.0001. Non-significant results are reported as ns.

**Fig. S5. Flow cytometry gating strategy for immune cell profiling panels.** Single cells isolated from tumor or inguinal lymph node tissues were stained with an antibody cocktail containing Zombie NIR, anti-CD45, anti-CD3, anti-CD19, anti-NKp46, anti-CD11b, anti-I-A/I-E, anti-Ly6G, anti-Ly6C, and anti-F4/80 antibodies. Doublets were excluded using FSC-H versus FSC-A; dead cells were excluded by gating out Zombie NIR-positive events; and immune cells were identified as the CD45^+^ population. Within the CD45^+^ population, T cells were identified by anti-CD3 antibody staining. From the CD3^−^ population, NK cells and B cells were identified using anti-NKp46 and anti-CD19 antibodies, respectively. The CD3^−^CD19^−^NKp46^−^ population was then gated on CD11b^+^ and I-A/I-E^+^ to identify the following populations: type 1 dendritic cells (CD11b^−^I-A/I-E^+^Ly-6G^−^Ly-6C^−^), type 2 dendritic cells (CD11b^+^I-A/I-E^+^Ly-6G^−^Ly-6C^−^), macrophages (CD11b^+^I-A/I-E+F4/80^+^), mMDSCs (CD11b^−^I-A/I-E^−^Ly-6G^−^Ly-6C^+^), and gMDSCs (CD11b^−^I-A/I-E^−^ Ly-6G^+^Ly-6C^−^). Black arrows indicate gated populations, and red arrows show identified immune cell populations.

**Fig. S6. Immune cell profiling of tumors and TDLNs from *Ulk1*-knockout mice reveals a reduction in total immune cell counts.** Immune cell profiling of tumor tissues from C57BL/6 mice injected with mEERL/sg-Scr or mEERL/sg2-*Ulk1* (n=4/group). Isolated single cell suspensions were stained with an antibody cocktail and analyzed by flow cytometry. Representative plots of CD45^+^ cells (A). Pie charts show the immune distribution of 8 types within the CD45^+^ population (B). Representative plots showing percentage of macrophages (C), NK cells (D), B cells (E), type 2 dendritic cells (F), type 1 Dendritic cells (G), monocytic MDSCs (H), and granulocytic MDSCs (I) to total CD45^+^ cells. *P-*value was determined by Student’s t-test. \**p* < 0.05, \*\**p* < 0.01. Non-significant results are reported as ns. T cell profiling of tumor draining lymph nodes (TDLNs) from mice injected with either mEERL/sg-Scr or mEERL/sg-*Ulk1* was isolated 21 days post-injection (n=3/group). Single-cell suspensions were stained with an antibody cocktail containing Zombie NIR, anti-CD45, anti-CD3, anti-CD4, anti-CD8, anti-CD44, and anti-CD62L antibodies. Representative histograms showing percentages of total T cells (J) and different effector states based on CD62L and CD44 (K).

## REFERENCES

1. Chaturvedi, A. K. et al. Human Papillomavirus and Rising Oropharyngeal Cancer Incidence in the United States. J. Clin. Oncol. 41, 3081–3088 (2023).

2. Gillison, M. L., Chaturvedi, A. K., Anderson, W. F. & Fakhry, C. Epidemiology of Human Papillomavirus-Positive Head and Neck Squamous Cell Carcinoma. J. Clin. Oncol. Off. J. Am. Soc. Clin. Oncol. 33, 3235–3242 (2015).

3. Piersma, S. J. Immunosuppressive Tumor Microenvironment in Cervical Cancer Patients. Cancer Microenviron. 4, 361–375 (2011).

4. Warren, C. J. et al. APOBEC3A Functions as a Restriction Factor of Human Papillomavirus. J. Virol. 89, 688–702 (2014).

5. Faden, D. L. et al. APOBEC mutagenesis is tightly linked to the immune landscape and immunotherapy biomarkers in head and neck squamous cell carcinoma. Oral Oncol. 96, 140–147 (2019).

6. Kansy, B. A. et al. HPV-associated head and neck cancer is characterized by distinct profiles of CD8+ T cells and myeloid-derived suppressor cells. Cancer Immunol. Immunother. CII 72, 4367–4383 (2023).

7. Kirchner, J. et al. Type I conventional dendritic cells and CD8+ T cells predict favorable clinical outcome of head and neck squamous cell carcinoma patients. Front. Immunol. 15, 1414298 (2024).

8. Hewavisenti, R. et al. CD103+ tumor-resident CD8+ T cell numbers underlie improved patient survival in oropharyngeal squamous cell carcinoma. J. Immunother. Cancer 8, e000452 (2020).

9. Näsman, A. et al. Tumor Infiltrating CD8+ and Foxp3+ Lymphocytes Correlate to Clinical Outcome and Human Papillomavirus (HPV) Status in Tonsillar Cancer. PLoS ONE 7, e38711 (2012).

10. Ferris, R. L. et al. Nivolumab for Recurrent Squamous-Cell Carcinoma of the Head and Neck. N. Engl. J. Med. 375, 1856–1867 (2016).

11. Cohen, E. E. W. et al. Pembrolizumab versus methotrexate, docetaxel, or cetuximab for recurrent or metastatic head-and-neck squamous cell carcinoma (KEYNOTE-040): a randomised, open-label, phase 3 study. The Lancet 393, 156–167 (2019).

12. Ugurel, S. et al. MHC class-I downregulation in PD-1/PD-L1 inhibitor refractory Merkel cell carcinoma and its potential reversal by histone deacetylase inhibition: a case series. Cancer Immunol. Immunother. CII 68, 983–990 (2019).

13. Montesion, M. et al. Somatic HLA Class I Loss Is a Widespread Mechanism of Immune Evasion Which Refines the Use of Tumor Mutational Burden as a Biomarker of Checkpoint Inhibitor Response. Cancer Discov. 11, 282–292 (2021).

14. D’Amico, S. et al. Targeting the antigen processing and presentation pathway to overcome resistance to immune checkpoint therapy. Front. Immunol. 13, 948297 (2022).

15. Khalil, M. I. et al. The membrane-associated ubiquitin ligase MARCHF8 promotes cancer immune evasion by degrading MHC class I proteins. Preprint at 10.1101/2024.11.29.626106 (2024) *PNAS* in press.

16. Goto, E. et al. c-MIR, a Human E3 Ubiquitin Ligase, Is a Functional Homolog of Herpesvirus Proteins MIR1 and MIR2 and Has Similar Activity. J. Biol. Chem. 278, 14657–14668 (2003).

17. Khalil, M. I. et al. HPV upregulates MARCHF8 ubiquitin ligase and inhibits apoptosis by degrading the death receptors in head and neck cancer. PLOS Pathog. 19, e1011171 (2023).

18. Bartee, E., Mansouri, M., Hovey Nerenberg, B. T., Gouveia, K. & Früh, K. Downregulation of Major Histocompatibility Complex Class I by Human Ubiquitin Ligases Related to Viral Immune Evasion Proteins. J. Virol. 78, 1109–1120 (2004).

19. Lehner, P. J., Hoer, S., Dodd, R. & Duncan, L. M. Downregulation of cell surface receptors by the K3 family of viral and cellular ubiquitin E3 ligases. Immunol. Rev. 207, 112–125 (2005).

20. Vargas, J. N. S., Hamasaki, M., Kawabata, T., Youle, R. J. & Yoshimori, T. The mechanisms and roles of selective autophagy in mammals. Nat. Rev. Mol. Cell Biol. 24, 167–185 (2023).

21. Autophagy in yeast demonstrated with proteinase-deficient mutants and conditions for its induction. J. Cell Biol. 119, 301–311 (1992).

22. Zaffagnini, G. & Martens, S. Mechanisms of Selective Autophagy. J. Mol. Biol. 428, 1714–1724 (2016).

23. Birgisdottir, Å. B., Lamark, T. & Johansen, T. The LIR motif – crucial for selective autophagy. J. Cell Sci. 126, 3237–3247 (2013).

24. Germain, K. et al. Upregulated pexophagy limits the capacity of selective autophagy. Nat. Commun. 15, 375 (2024).

25. Mancias, J. D. & Kimmelman, A. C. Mechanisms of selective autophagy in normal physiology and cancer. J. Mol. Biol. 428, 1659–1680 (2016).

26. Adriaenssens, E., Ferrari, L. & Martens, S. Orchestration of selective autophagy by cargo receptors. Curr. Biol. 32, R1357–R1371 (2022).

27. Fujita, N. et al. The Atg16L Complex Specifies the Site of LC3 Lipidation for Membrane Biogenesis in Autophagy. Mol. Biol. Cell 19, 2092–2100 (2008).

28. Chang, C. et al. Reconstitution of cargo-induced LC3 lipidation in mammalian selective autophagy. Sci. Adv. 7, eabg4922 (2021).

29. Nath, S. et al. Lipidation of the LC3/GABARAP family of autophagy proteins relies upon a membrane curvature-sensing domain in Atg3. Nat. Cell Biol. 16, 415–424 (2014).

30. Yang, Y. et al. Silencing of LncRNA-HOTAIR decreases drug resistance of Non-Small Cell Lung Cancer cells by inactivating autophagy via suppressing the phosphorylation of ULK1. Biochem. Biophys. Res. Commun. 497, 1003–1010 (2018).

31. Zhan, L., Zhang, J., Wei, B. & Cao, Y. Selective autophagy of NLRC5 promotes immune evasion of endometrial cancer. Autophagy 18, 942–943.

32. Jin, Y., Qiu, J., Lu, X. & Li, G. C-MYC Inhibited Ferroptosis and Promoted Immune Evasion in Ovarian Cancer Cells through NCOA4 Mediated Ferritin Autophagy. Cells 11, 4127 (2022).

33. Yamamoto, K. et al. Autophagy promotes immune evasion of pancreatic cancer by degrading MHC-I. Nature 581, 100–105 (2020).

34. Pyeon, D. et al. Fundamental Differences in Cell Cycle Deregulation in Human Papillomavirus–Positive and Human Papillomavirus–Negative Head/Neck and Cervical Cancers. Cancer Res. 67, 4605–4619 (2007).

35. Gameiro, S. F. et al. Analysis of Class I Major Histocompatibility Complex Gene Transcription in Human Tumors Caused by Human Papillomavirus Infection. Viruses 9, 252 (2017).

36. Doench, J. G. et al. Optimized sgRNA design to maximize activity and minimize off-target effects of CRISPR-Cas9. Nat. Biotechnol. 34, 184–191 (2016).

37. Li, W. et al. MAGeCK enables robust identification of essential genes from genome-scale CRISPR/Cas9 knockout screens. Genome Biol. 15, 554 (2014).

38. Brinkman, E. K. & van Steensel, B. Rapid Quantitative Evaluation of CRISPR Genome Editing by TIDE and TIDER. in CRISPR Gene Editing: Methods and Protocols (ed. Luo, Y.) 29–44 (Springer, New York, NY, 2019). doi:10.1007/978-1-4939-9170-9_3.

39. Chakrabarti, A. M. et al. Target-Specific Precision of CRISPR-Mediated Genome Editing. Mol. Cell 73, 699–713.e6 (2019).

40. Sentmanat, M. F., Peters, S. T., Florian, C. P., Connelly, J. P. & Pruett-Miller, S. M. A Survey of Validation Strategies for CRISPR-Cas9 Editing. Sci. Rep. 8, 888 (2018).

41. Brinkman, E. K., Chen, T., Amendola, M. & van Steensel, B. Easy quantitative assessment of genome editing by sequence trace decomposition. Nucleic Acids Res. 42, e168 (2014).

42. Russell, R. C. et al. ULK1 induces autophagy by phosphorylating Beclin-1 and activating Vps34 lipid kinase. Nat. Cell Biol. 15, 741–750 (2013).

43. Loos, B., du Toit, A. & Hofmeyr, J.-H. S. Defining and measuring autophagosome flux—concept and reality. Autophagy 10, 2087–2096 (2014).

44. Martin, K. R. et al. A Potent and Selective ULK1 Inhibitor Suppresses Autophagy and Sensitizes Cancer Cells to Nutrient Stress. iScience 8, 74–84 (2018).

45. Nagasaki, J., Ishino, T. & Togashi, Y. Mechanisms of resistance to immune checkpoint inhibitors. Cancer Sci. 113, 3303–3312 (2022).

46. Gu, S. S. et al. Therapeutically increasing MHC-I expression potentiates immune checkpoint blockade. Cancer Discov. 11, 1524–1541 (2021).

47. Sade-Feldman, M. et al. Resistance to checkpoint blockade therapy through inactivation of antigen presentation. Nat. Commun. 8, 1136 (2017).

48. Pembrolizumab versus methotrexate, docetaxel, or cetuximab for recurrent or metastatic head-and-neck squamous cell carcinoma (KEYNOTE-040): a randomised, open-label, phase 3 study - ScienceDirect. https://www.sciencedirect.com/science/article/pii/S0140673618319998?via%3Dihub.

49. Kansy, B. A. et al. PD-1 status in CD8+ T cells associates with survival and anti-PD-1 therapeutic outcomes in head and neck cancer. Cancer Res. 77, 6353–6364 (2017).

50. Welters, M. J. P. et al. Intratumoral HPV16-Specific T Cells Constitute a Type I–Oriented Tumor Microenvironment to Improve Survival in HPV16-Driven Oropharyngeal Cancer. Clin. Cancer Res. 24, 634–647 (2018).

51. Herhaus, L. et al. IRGQ-mediated autophagy in MHC class I quality control promotes tumor immune evasion. Cell S0092867424011486 (2024) doi:10.1016/j.cell.2024.09.048.

52. Sang, W. et al. Receptor-interacting Protein Kinase 2 Is an Immunotherapy Target in Pancreatic Cancer. Cancer Discov. 14, 326–347 (2024).

53. Chen, X. et al. A membrane-associated MHC-I inhibitory axis for cancer immune evasion. Cell 186, 3903–3920.e21 (2023).

54. Young, T. M. et al. Autophagy protects tumors from T cell–mediated cytotoxicity via inhibition of TNFα-induced apoptosis. Sci. Immunol. 5, eabb9561 (2020).

55. Orchestration of selective autophagy by cargo receptors - ScienceDirect. https://www.sciencedirect.com/science/article/pii/S0960982222017584?via%3Dihub.

56. Ravenhill, B. J. et al. The Cargo Receptor NDP52 Initiates Selective Autophagy by Recruiting the ULK Complex to Cytosol-Invading Bacteria. Mol. Cell 74, 320–329.e6 (2019).

57. Patil, C. D. & Shukla, D. OPTN (optineurin)-mediated selective autophagy prevents neurodegeneration due to herpesvirus infection. Autophagy 18, 944–945.

58. Xie, X. et al. Molecular basis of ubiquitin recognition by the autophagy receptor CALCOCO2. Autophagy 11, 1775–1789 (2015).

59. Cemma, M., Kim, P. K. & Brumell, J. H. The ubiquitin-binding adaptor proteins p62/SQSTM1 and NDP52 are recruited independently to bacteria-associated microdomains to target Salmonella to the autophagy pathway. Autophagy 7, 341–345 (2011).

60. Mostowy, S. et al. p62 and NDP52 Proteins Target Intracytosolic Shigella and Listeria to Different Autophagy Pathways. J. Biol. Chem. 286, 26987–26995 (2011).

61. Tracz, M. & Bialek, W. Beyond K48 and K63: non-canonical protein ubiquitination. Cell. Mol. Biol. Lett. 26, 1 (2021).

62. Manohar, S. et al. Polyubiquitin Chains Linked by Lysine Residue 48 (K48) Selectively Target Oxidized Proteins In Vivo. Antioxid. Redox Signal. 31, 1133–1149 (2019).

63. Chen, Z. et al. Duck MARCH8 Negatively Regulates the RLR Signaling Pathway through K29-Linked Polyubiquitination of MAVS. J. Immunol. 210, 786–794 (2023).

64. Jin, S. et al. Tetherin Suppresses Type I Interferon Signaling by Targeting MAVS for NDP52-Mediated Selective Autophagic Degradation in Human Cells. Mol. Cell 68, 308–322.e4 (2017).

65. Loi, M. et al. Macroautophagy Proteins Control MHC Class I Levels on Dendritic Cells and Shape Anti-viral CD8+ T Cell Responses. Cell Rep. 15, 1076–1087 (2016).

66. Øynebråten, I. Involvement of autophagy in MHC class I antigen presentation. Scand. J. Immunol. 92, e12978 (2020).

67. Yu, C. et al. MARCH8 Inhibits Ebola Virus Glycoprotein, Human Immunodeficiency Virus Type 1 Envelope Glycoprotein, and Avian Influenza Virus H5N1 Hemagglutinin Maturation. mBio 11, e01882-20 (2020).

68. Sankar, D. S. & Dengjel, J. Interactors and neighbors of ULK1 complex members. Autophagy 21, 243–245.

69. Sankar, D. S. et al. The ULK1 effector BAG2 regulates autophagy initiation by modulating AMBRA1 localization. Cell Rep. 43, 114689 (2024).

70. Lee, E.-J. & Tournier, C. The requirement of uncoordinated 51-like kinase 1 (ULK1) and ULK2 in the regulation of autophagy. Autophagy 7, 689–695 (2011).

71. Cheong, H., Lindsten, T., Wu, J., Lu, C. & Thompson, C. B. Ammonia-induced autophagy is independent of ULK1/ULK2 kinases. Proc. Natl. Acad. Sci. U. S. A. 108, 11121–11126 (2011).

72. Demeter, A. et al. ULK1 and ULK2 are less redundant than previously thought: computational analysis uncovers distinct regulation and functions of these autophagy induction proteins. Sci. Rep. 10, 10940 (2020).

73. Li, X. et al. ULK2 promotes migration and invasion of colorectal cancer cells via MCT4-mediated lactate export. Med. Oncol. 42, 368 (2025).

74. Vaughan, R. M., Dickson, B. M., Martin, K. R. & MacKeigan, J. P. Molecular dynamics simulations provide insights into ULK-101 potency and selectivity toward autophagic kinases ULK1/2. J. Biomol. Struct. Dyn. 43, 7106–7113 (2025).

75. Noman, M. Z. et al. Inhibition of Vps34 reprograms cold into hot inflamed tumors and improves anti–PD-1/PD-L1 immunotherapy. Sci. Adv. 6, eaax7881 (2020).

76. Mgrditchian, T. et al. Targeting autophagy inhibits melanoma growth by enhancing NK cells infiltration in a CCL5-dependent manner. Proc. Natl. Acad. Sci. U. S. A. 114, E9271–E9279 (2017).

77. Callejas-Valera, J. L. et al. Characterization of the Immune Response to PD-1 Blockade during Chemoradiotherapy for Head and Neck Squamous Cell Carcinoma. Cancers 14, 2499 (2022).

78. Griffin, L. M., Cicchini, L. & Pyeon, D. Human papillomavirus infection is inhibited by host autophagy in primary human keratinocytes. Virology 437, 12–19 (2013).

79. Kiritsy, M. C. et al. A genetic screen in macrophages identifies new regulators of IFNγ-inducible MHCII that contribute to T cell activation. eLife 10, e65110.

80. Labun, K. et al. CHOPCHOP v3: expanding the CRISPR web toolbox beyond genome editing. Nucleic Acids Res. 47, W171–W174 (2019).

81. Brinkman, E. K. et al. Easy quantification of template-directed CRISPR/Cas9 editing. Nucleic Acids Res. 46, e58 (2018).

82. Iwabuchi, S., Kakazu, Y., Koh, J.-Y. & Charles Harata, N. Evaluation of the effectiveness of Gaussian filtering in distinguishing punctate synaptic signals from background noise during image analysis. J. Neurosci. Methods 223, 92–113 (2014).

