## Supplementary figures and images for "ULK1 drives NDP52-mediated selective autophagic degradation of MHC-I to promote immune evasion in HPV-positive head and neck cancer"

### Supplementary Figures

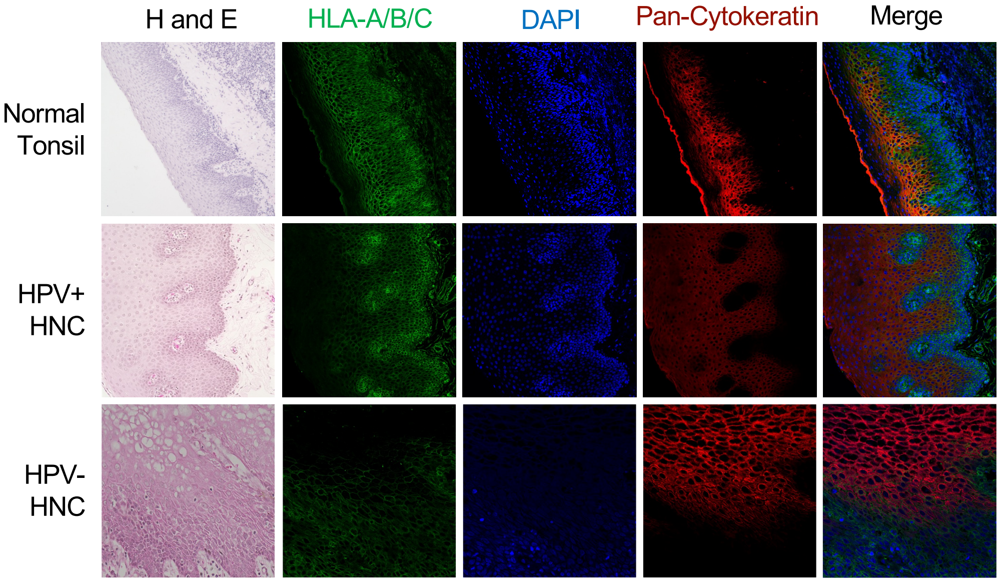

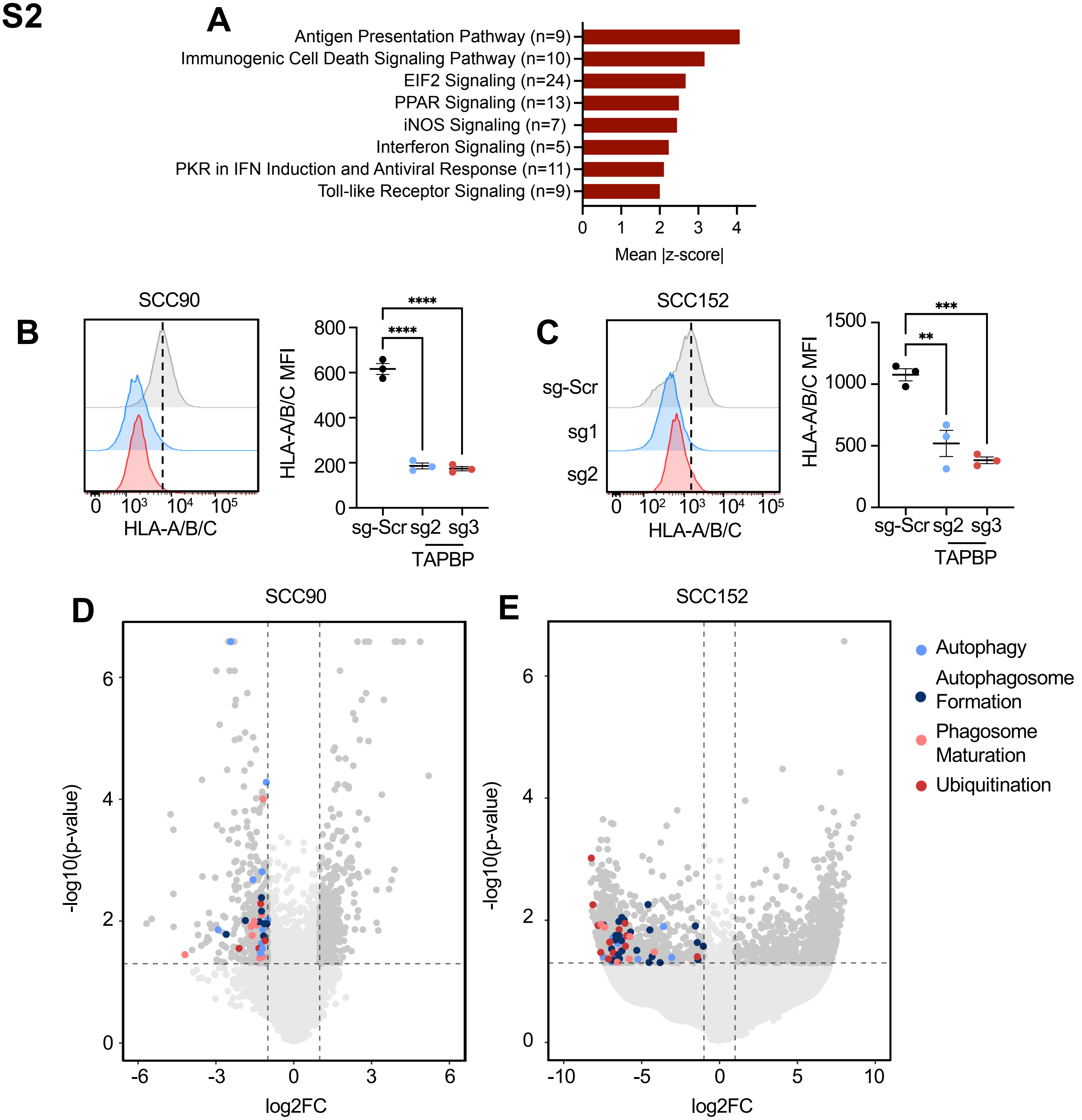

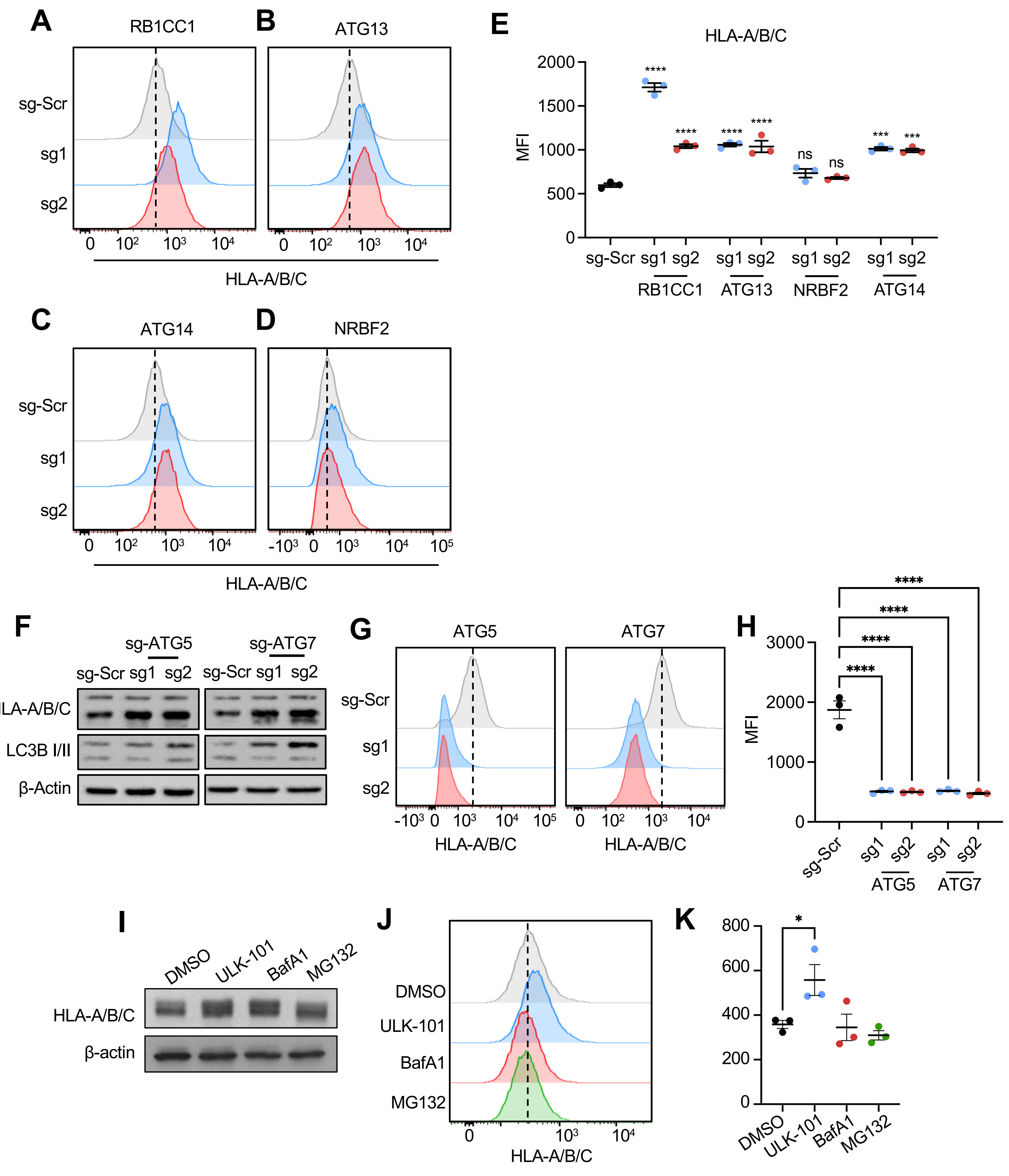

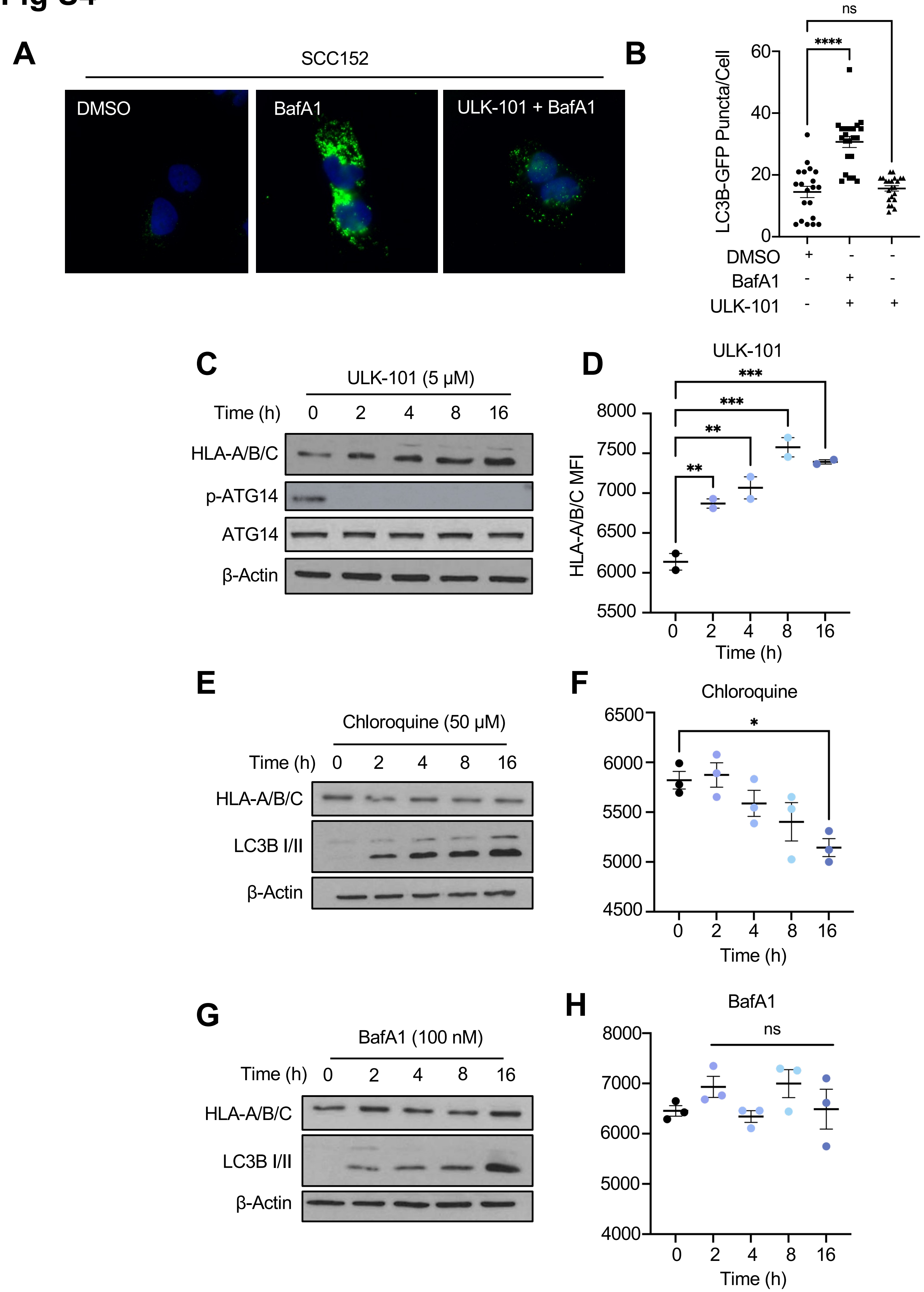

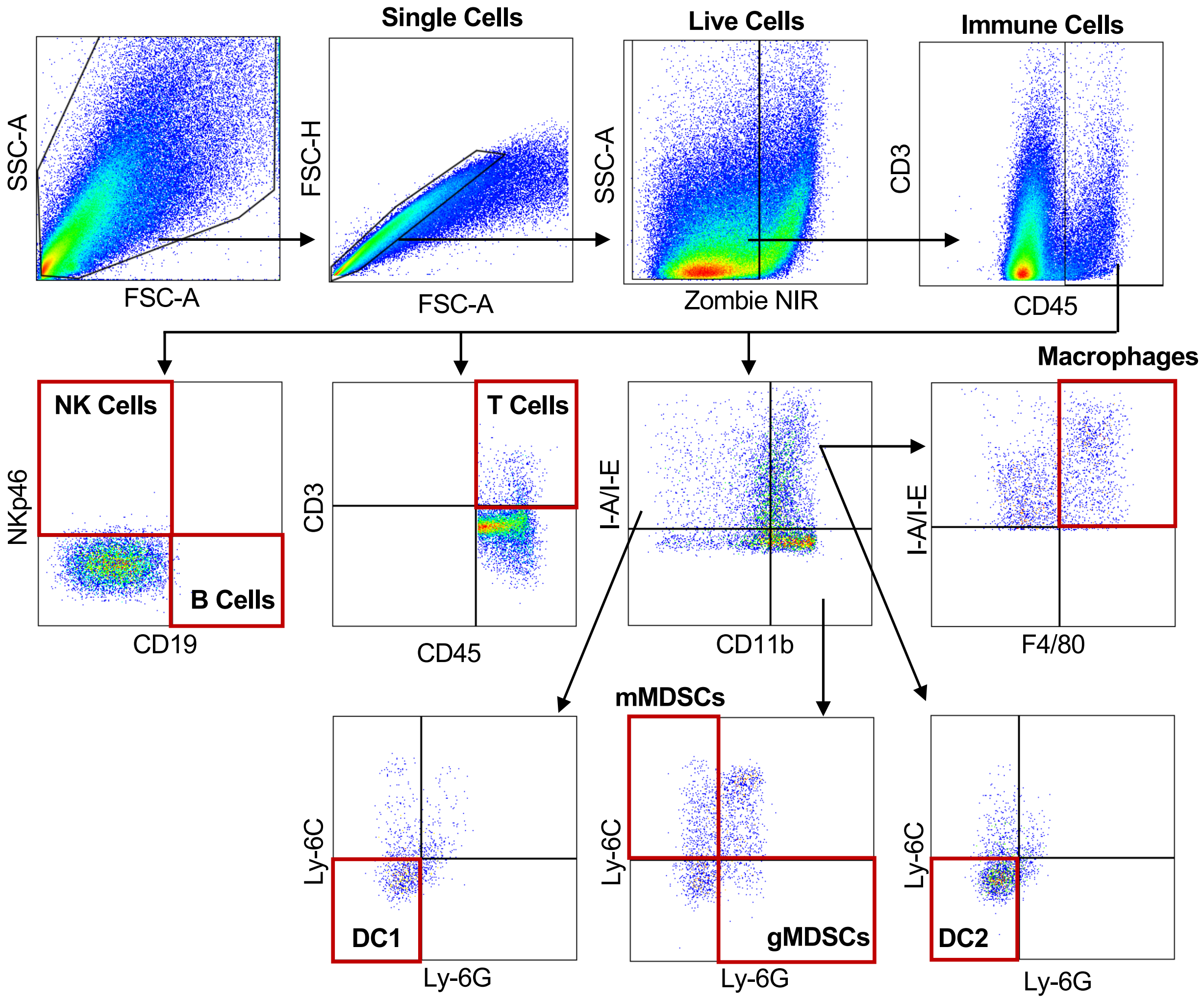

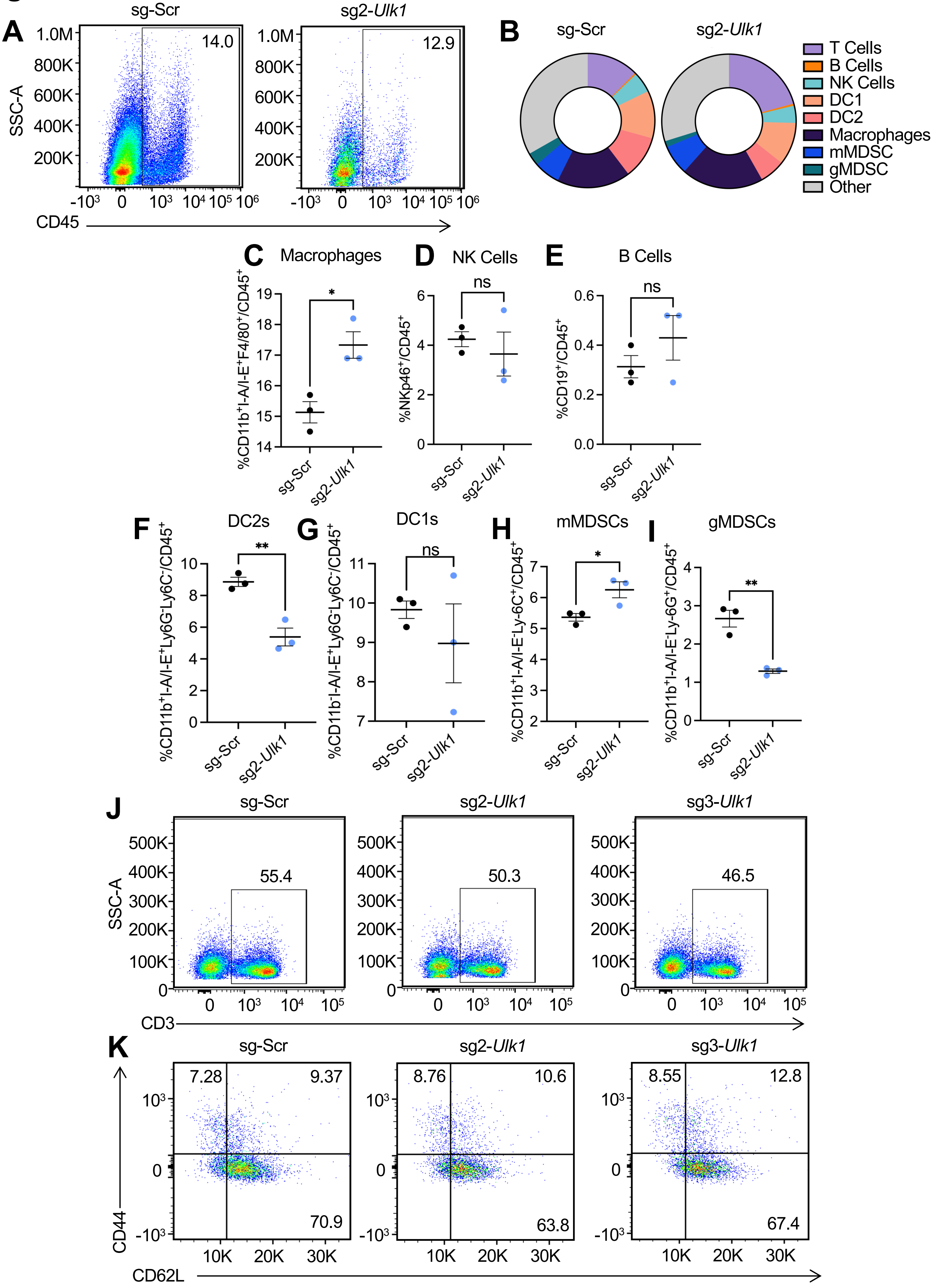
