## Supplementary Tables for "ULK1 drives NDP52-mediated selective autophagic degradation of MHC-I to promote immune evasion in HPV-positive head and neck cancer"

**Table S1. Patient tumor sample information.**

| Sample ID | HPV Status | Site | AJCC Edition | TNM Stage | Stage | Procedure | Treatment |
|---|---|---|---|---|---|---|---|
| DCA 019-1048 | HPV positive | Tonsil | 7 | cT2N2b, pT2NX | IVa | Resection | Untreated |
| DCA 022-1035 | HPV positive | Tonsil | 7 | cT2N1 | III | Biopsy | Untreated |
| DCA 029-1250 | HPV positive | Tonsil | 7 | cT3N2b, pT2NX | IVa | Resection | Untreated |
| DCA 031-1296 | HPV positive | Tonsil | 7 | cT2N2b, pT2NX | IVa | Resection | Untreated |
| DCA 036-1223 | HPV positive | Tonsil | 7 | cT2N2c | IVa | Biopsy | Untreated |
| DCA 038-1165 | HPV positive | BOT | 7 | cT1N2a | IVa | Biopsy | Untreated |
| DCA 043-1253 | HPV positive | Tonsil | 7 | cT4aN2b | IVa | Biopsy | Untreated |
| DCA FG 1124 | HPV positive | Tonsil | 7 | cT1N2a, pT1NX | IVa | Biopsy | Untreated |
| DCA FG 1140 | HPV positive | Tonsil | 7 | cT1N2a, pT1NX | IVa | Biopsy | Untreated |
| HN 006 | HPV positive | BOT | 6 | cT3N2b | IVa | Biopsy | Untreated |
| HN 011 | HPV positive | Tonsil | 6 | cT2N2b | IVa | Biopsy | Untreated |
| HN 017 | HPV positive | R. tonsil | 6 | cT2N2b, pT2N2b | IVa | Salvage neck dissection | Post-CRT |
| HN 033 | HPV positive | Oropharynx | 6 | cT4N2c | IVa | Biopsy | Untreated* |
| HN 056 | HPV positive | BOT | 7 | cT2N2c | IVa | Biopsy | Untreated |
| HN 061 | HPV positive | R. tonsil | 7 | cT2N2a | IVa | Biopsy | Untreated |
| HN 062 | HPV positive | BOT | 7 | cT4N2c | IVa | Biopsy | Untreated |
| HN 064 | HPV positive | R. tonsil | 7 | cT1N2b, pT3N2b | IVa | Biopsy | Untreated |
| HN 065 | HPV positive | Tonsil | 7 | cT4bN2c | IVb | Biopsy | Untreated |
| HN 084 | HPV positive | Tonsil | 7 | cT2N2b, pT2N2b | IVa | Resection | Untreated |
| HN 094 | HPV positive | Tonsil | 7 | cT2N2b | IVa | Biopsy | Untreated |
| HN 121 | HPV positive | Tonsillar pillar | 7 | cT1N1, pT1N0 | I | Resection | Untreated |
| HN 161 | HPV positive | Oropharynx | 7 | cT2N3 | IVb | Biopsy | Untreated |
| HN 370 | HPV positive | BOT | 7 | cT1N1, pT1NX | III | Resection | Untreated |
| DCA 015-1237 | HPV negative | Tonsil | 7 | cT4aN1 | IVa | Biopsy | Untreated |
| HN 003 | HPV negative | R. tonsil | 6 | cT2N2b | IVa | Biopsy | Untreated |
| HN 013 | HPV negative | Oropharynx | 6 | cT1N0 | I | Biopsy | Untreated |
| HN 026 | HPV negative | Oropharynx | 6 | cT3N2b | IVa | Resection | Untreated |
| HN 053 | HPV negative | R. tonsil | 7 | cT1N2c, pT1N2a | IVa | Resection | Untreated |
| HN 057 | HPV negative | R. tonsil | 7 | cT4aN2b, pT4aN2b | IVa | Biopsy | Untreated |
| HN 169 | HPV negative | R. tonsil | 7 | cT1N0, pT1N0 | I | Resection | Untreated |
| HN 185 | HPV negative | Tonsil | 7 | cT2N0, pT1NX | II | Resection | Untreated |
| HN 232 | HPV negative | Tonsil | 7 | cT1N2b, pT2N2b | IVa | Resection | Untreated |
| HN 235 | HPV negative | BOT | 7 | cT1N2b, pT1N2b | IVa | Resection | Untreated |
| HN 253 | HPV negative | BOT, FOM | 7 | cT2N0, pT2NX | II | Resection | Untreated |
| HN 310 | HPV negative | Tonsillar pillar | 7 | cT2N2b, pT1N2b | IVa | Resection | Untreated |
| 09-26815 | Normal tonsil | Tonsil | | | | | |

| | | |
|---|---|---|
| 19x38734 | Normal tonsil | Tonsil |
| 10-11318 | Normal tonsil | Tonsil |
| 09-25936 | Normal tonsil | Tonsil |
| 18x05001 | Normal tonsil | Tonsil |
| 18x40899 | Normal tonsil | Tonsil |
| 19x40087 | Normal tonsil | Tonsil |
| 19x45743 | Normal tonsil | Tonsil |
| 19x40646 | Normal tonsil | Tonsil |
| 19x40648 | Normal tonsil | Tonsil |
| 19x46251 | Normal tonsil | Tonsil |
| 19x44612 | Normal tonsil | Tonsil |

AJCC, American Joint Committee on Cancer; TNM, Tumor Node Metastasis; cTNM, clinical staging; pTNM, pathologic staging; BOT, Base of tongue; R. Tonsil, Right Tonsil; FOM, Floor of Mouth; CRT, chemoradiotherapy

**Table S2. CRISPR screen library preparation primers.**

| Name | Sequence 5'→3' |
|---|---|
| P5 0 nt stagger | AATGATACGGCGACCACCGAGATCTACACTCTTTCCCTACACGACGCTCTTCCGATCTTTGTGGAAAGGACGAAACACCG |
| P5 1nt stagger | AATGATACGGCGACCACCGAGATCTACACTCTTTCCCTACACGACGCTCTTCCGATCTCTTGTGGAAAGGACGAAACACCG |
| P5 2 nt stagger | AATGATACGGCGACCACCGAGATCTACACTCTTTCCCTACACGACGCTCTTCCGATCTGCTTGTGGAAAGGACGAAACACCG |
| P5 3 nt stagger | AATGATACGGCGACCACCGAGATCTACACTCTTTCCCTACACGACGCTCTTCCGATCTAGCTTGTGGAAAGGACGAAACACCG |
| P5 4 nt stagger | AATGATACGGCGACCACCGAGATCTACACTCTTTCCCTACACGACGCTCTTCCGATCTCAACTTGTGGAAAGGACGAAACACCG |
| P5 6 nt stagger | AATGATACGGCGACCACCGAGATCTACACTCTTTCCCTACACGACGCTCTTCCGATCTTGCACCTTGTGGAAAGGACGAAACACCG |
| P5 7nt stagger | AATGATACGGCGACCACCGAGATCTACACTCTTTCCCTACACGACGCTCTTCCGATCTACGCAACTTGTGGAAAGGACGAAACACCG |
| P5 8nt stagger | AATGATACGGCGACCACCGAGATCTACACTCTTTCCCTACACGACGCTCTTCCGATCTGAAGACCCTTGTGGAAAGGACGAAACACCG |
| P7_D701 | CAAGCAGAAGACGGCATACGAGATATTACTCGGTGACTGGAGTTCAGACGTGTGCTCTTCCGATCTTCTACTATTCTTTCCCCTGCACTGT |
| P7_D702 | CAAGCAGAAGACGGCATACGAGATTCCGGAGAGTGACTGGAGTTCAGACGTGTGCTCTTCCGATCTTCTACTATTCTTTCCCCTGCACTGT |
| P7_D703 | CAAGCAGAAGACGGCATACGAGATCGCTCATTGTGACTGGAGTTCAGACGTGTGCTCTTCCGATCTTCTACTATTCTTTCCCCTGCACTGT |

| | |
|---|---|
| P7_D704 | CAAGCAGAAGACGGCATACGAGATGAGATTCCGTGACTGGAGTTCAGACGTGTGCTCTTCCGATCTTCTACTATTCTTTCCCCTGCACTGT |
| P7_D705 | CAAGCAGAAGACGGCATACGAGATATTCAGAAGTGACTGGAGTTCAGACGTGTGCTCTTCCGATCTTCTACTATTCTTTCCCCTGCACTGT |
| P7_D706 | CAAGCAGAAGACGGCATACGAGATGAATTCGTGTGACTGGAGTTCAGACGTGTGCTCTTCCGATCTTCTACTATTCTTTCCCCTGCACTGT |
| P7_D707 | CAAGCAGAAGACGGCATACGAGATCTGAAGCTGTGACTGGAGTTCAGACGTGTGCTCTTCCGATCTTCTACTATTCTTTCCCCTGCACTGT |
| P7_D708 | CAAGCAGAAGACGGCATACGAGATTAATGCGCGTGACTGGAGTTCAGACGTGTGCTCTTCCGATCTTCTACTATTCTTTCCCCTGCACTGT |
| P7_D709 | CAAGCAGAAGACGGCATACGAGATCGGCTATGGTGACTGGAGTTCAGACGTGTGCTCTTCCGATCTTCTACTATTCTTTCCCCTGCACTGT |
| P7_D710 | CAAGCAGAAGACGGCATACGAGATTCCGCGAAGTGACTGGAGTTCAGACGTGTGCTCTTCCGATCTTCTACTATTCTTTCCCCTGCACTGT |
| P7_D711 | CAAGCAGAAGACGGCATACGAGATTCTCGCGCGTGACTGGAGTTCAGACGTGTGCTCTTCCGATCTTCTACTATTCTTTCCCCTGCACTGT |
| P7_D712 | CAAGCAGAAGACGGCATACGAGATAGCGATAGGTGACTGGAGTTCAGACGTGTGCTCTTCCGATCTTCTACTATTCTTTCCCCTGCACTGT |

nt, nucleotide

**Table S3. Top gene hits from CRISPR screens.**

| Cell Line | Pathway | Gene | p-value | Log2FC |
|---|---|---|---|---|
| SCC90 | Autophagy | DAPK2 | 0.013896 | -2.9161 |
| | | PRKAG3 | 0.0095392 | -1.0037 |
| | | PPP2R1A | 0.023097 | -1.2411 |
| | | PPP2R2A | 0.0015497 | -1.2247 |
| | | STK11 | 0.013681 | -1.19 |
| | | ATG101 | 5.26E-05 | -1.0583 |
| | | ATG4D | 0.0020869 | -1.5723 |
| | | SEC62 | 0.028191 | -1.2075 |
| | | TMEM127 | 2.59E-07 | -2.4303 |
| | | WWP2 | 0.033449 | -1.2832 |
| | Autophagosome Formation | ARPC4 | 0.017569 | -1.1626 |
| | | FFAR1 | 0.011026 | -1.0298 |
| | | GNRHR | 0.016507 | -2.6035 |
| | | PIP4K2C | 0.011034 | -1.138 |
| | | PLA2G4C | 0.0041237 | -1.2444 |
| | | PTGER4 | 0.0097847 | -1.8671 |
| | | SLC52A1 | 0.0068589 | -1.2422 |
| | | TLR7 | 0.010218 | -1.3321 |
| | Phagosome Maturation | VPS16 | 0.04047 | -1.3148 |
| | | VPS33A | 9.92E-05 | -1.1801 |
| | | ATP6V0C | 0.0172 | -1.6021 |
| | | ATP6V1B2 | 0.035492 | -4.1933 |
| | | ATP6V1F | 0.03834 | -1.1905 |
| | | ATP6V1G1 | 0.012223 | -1.6496 |
| | | GPAA1 | 0.011542 | -1.4366 |
| | | PRDX1 | 0.0079084 | -1.2382 |
| | | VAMP2 | 0.010068 | -1.5164 |
| | Ubiquitination | DNAJC1 | 0.0051675 | -1.2736 |
| | | PSMD1 | 0.027994 | -2.1084 |
| | | STUB1 | 0.02808 | -1.344 |
| | | UBE2R2 | 0.020954 | -1.0998 |
| SCC152 | Autophagy | ATG2A | 0.040818 | -3.0706 |
| | | EGF | 0.048041 | -6.9005 |
| | | GNB1L | 0.012132 | -7.7967 |
| | | IRS1 | 0.02402 | -6.5304 |
| | | KAT5 | 0.019559 | -6.8592 |
| | | PPP2R3B | 0.012641 | -3.5924 |
| | | RB1CC1 | 0.043445 | -5.2359 |
| | | TNFRSF1A | 0.036252 | -6.5676 |
| | | ULK1 | 0.039791 | -7.4647 |
| | Autophagosome Formation | ADGRB1 | 0.039301 | -4.3287 |
| | | ADGRF1 | 0.01049 | -6.4615 |
| | | ADORA2B | 0.02659 | -1.042 |
| | | BDKRB1 | 0.044171 | -1.3841 |
| | | CMKLR1 | 0.0055335 | -4.5932 |
| | | EDNRA | 0.012393 | -1.5534 |
| | | ERAS | 0.014424 | -4.4714 |

| | | | |
|---|---|---|---|
| | FCER2 | 0.015581 | -5.709 |
| | GCGR | 0.048985 | -4.5256 |
| | GRM5 | 0.023112 | -1.4277 |
| | HRH3 | 0.011973 | -7.4434 |
| | HTR1B | 0.017706 | -6.6624 |
| | HTR4 | 0.029348 | -6.9309 |
| | ITGAM | 0.031611 | -6.2632 |
| | MYH8 | 0.031114 | -5.3139 |
| | MYO18A | 0.042658 | -6.4235 |
| | NTSR2 | 0.021027 | -6.6841 |
| | PLA2G10 | 0.04484 | -6.8127 |
| | PLA2G3 | 0.0099765 | -6.1127 |
| | PNPLA2 | 0.021513 | -6.2992 |
| | TACR3 | 0.036679 | -6.4943 |
| | TAS1R1 | 0.049322 | -3.8199 |
| | TIMD4 | 0.0090075 | -6.2743 |
| | VIPR2 | 0.017774 | -6.4525 |
| Phagosome Maturation | ATP6V1G2 | 0.03302 | -4.1875 |
| | DYNC1I1 | 0.043059 | -5.7985 |
| | HLA-DRB1 | 0.031877 | -6.3915 |
| | MR1 | 0.021497 | -6.2588 |
| | NCF2 | 0.023211 | -7.0853 |
| | RAB5B | 0.012911 | -7.3885 |
| | TUBA8 | 0.018503 | -5.7581 |
| | VAMP3 | 0.04848 | -6.5655 |
| | VPS37B | 0.011603 | -7.6175 |
| Ubiquitination | HSPA4L | 0.039565 | -1.4248 |
| | PAN2 | 0.017905 | -5.9681 |
| | PSMC5 | 0.042869 | -7.1403 |
| | PSMD13 | 0.0055735 | -8.1315 |
| | PSMD14 | 0.034297 | -6.826 |
| | PSMD4 | 0.022724 | -7.0386 |
| | UBE3A | 0.00096061 | -8.2309 |
| | UCHL3 | 0.011234 | -6.0682 |
| | USP14 | 0.014213 | -6.4437 |
| | USP20 | 0.01219 | -7.7945 |
| | USP39 | 0.033652 | -7.6274 |
| | XIAP | 0.026537 | -6.0412 |

Log2FC, $Log_2$(Fold Change)

**Table S4. sgRNA oligos**

| Name | Sequence 5' → 3' |
|---|---|
| RB1CC1 sg1 Fwd | CACCGTTTCTAACAGCTCTATTACG |
| RB1CC1 sg1 Rev | AAACCGTAATAGAGCTGTTAGAAAC |
| RB1CC1 sg2 Fwd | CACCGCTGGTTAGGCACTCCAACAG |
| RB1CC1 sg2 Rev | AAACCTGTTGGAGTGCCTAACCAGC |
| ATG13 sg1 Fwd | CACCGTTTACCCAATCTGAACCCGT |

| | |
|---|---|
| ATG13 sg1 Rev | AAACACGGGTTCAGATTGGGTAAAC |
| ATG13 sg2 Fwd | CACCGATGTGAACTCACCTACTGGA |
| ATG13 sg2 Rev | AAACTCCAGTAGGTGAGTTCACATC |
| NRBF2 sg1 Fwd | CACCGCTCAGGCAGGCATTTCTCTG |
| NRBF2 sg1 Rev | AAACCAGAGAAATGCCTGCCTGAGC |
| NRBF2 sg2 Fwd | CACCGTGAGGAGGAGCTGTTTCATA |
| NRBF2 sg2 Rev | AAACTATGAAACAGCTCCTCCTCAC |
| PIK3C3 sg1 Fwd | CACCGATACACATCCCATATGGTGA |
| PIK3C3 sg1 Rev | AAACTCACCATATGGGATGTGTATC |
| PIK3C3 sg2 Fwd | CACCGTAACTTACCATAGACATCTG |
| PIK3C3 sg2 Rev | AAACCAGATGTCTATGGTAAGTTAC |
| ATG14 sg1 Fwd | CACCGAGGAAGTAAAGACGGGTGTG |
| ATG14 sg1 Rev | AAACCACACCCGTCTTTACTTCCTC |
| ATG14 sg2 Fwd | CACCGCAGCACTGATGGTGTAGGCA |
| ATG14 sg2 Rev | AAACTGCCTACACCATCAGTGCTGC |
| NDP52 sg1 Fwd | CACCGCAGCAGGAAGTCCAATTCAA |
| NDP52 sg1 Rev | AAACTTGAATTGGACTTCCTGCTGC |
| NDP52 sg2 Fwd | CACCGTCAGGTCATCTTTAACAGTG |
| NDP52 sg2 Rev | AAACCACTGTTAAAGATGACCTGAC |
| SQSTM1 sg1 Fwd | CACCGCCTCACCTGATTCTGCCGTG |
| SQSTM1 sg1 Rev | AAACCACGGCAGAATCAGGTGAGGC |
| SQSTM1 sg2 Fwd | CACCGTGGCTCCGGAAGGTGAAACA |
| SQSTM1 sg2 Rev | AAACTGTTTCACCTTCCGGAGCCAC |
| NBR1 sg1 Fwd | CACCGTCTGTGTCACATGGAACAAG |
| NBR1 sg1 Rev | AAACCTTGTTCCATGTGACACAGAC |
| NBR1 sg2 Fwd | CACCGATGATACTGCACCAGACCCG |
| NBR1 sg2 Rev | AAACCGGGTCTGGTGCAGTATCATC |
| ATG5 sg1 Fwd | CACCGTGATATAGCGTGAAACAAGT |
| ATG5 sg1 Rev | AAACACTTGTTTCACGCTATATCAC |
| ATG5 sg2 Fwd | CACCGCCTTAGATGGACAGTGCAGA |
| ATG5 sg2 Rev | AAACTCTGCACTGTCCATCTAAGGC |
| ATG7 sg1 Fwd | CACCGTCCTACTTTAGACTTGGACA |
| ATG7 sg1 Rev | AAACTGTCCAAGTCTAAAGTAGGAC |
| ATG7 sg2 Fwd | CACCGCTCTTGTAAATACCATCTGT |
| ATG7 sg2 Rev | AAACACAGATGGTATTTACAAGAGC |
| TAP1 sg1 Fwd | CACCGCATCATGTCTCGGGTAACAG |
| TAP1 sg1 Rev | AAACCTGTTACCCGAGACATGATGC |
| TAP1 sg2 Fwd | CACCGGGCTCCAAGAGCGAAAACGC |
| TAP1 sg2 Rev | AAACGCGTTTTCGCTCTTGGAGCCC |
| TAP2 sg1 Fwd | CACCGGTTGATTCGAGACATGGTGT |
| TAP2 sg1 Rev | AAACACACCATGTCTCGAATCAAC |
| TAP2 sg2 Fwd | CACCGATCCCCATATATGTATACCA |
| TAP2 sg2 Rev | AAACTGGTATACATATATGGGGATC |
| TAPBP sg1 Fwd | CACCGGATCGAGTGTTGGTTCGTGG |
| TAPBP sg1 Rev | AAACCCACGAACCAACACTCGATCC |
| TAPBP sg2 Fwd | CACCGAAGCGGCTCATCTCGCAGTG |
| TAPBP sg2 Rev | AAACCACTGCGAGATGAGCCGCTTC |
| Mouse Ulk1 sg2 Fwd | CACCGCGGCCCGCTGAAGACACCCG |
| Mouse Ulk1 sg2 Rev | AAACCGGGTGTCTTCAGCGGGCCGC |
| Mouse *Ulk1* sg3 Fwd | CACCGTAGTCTGCGTACCACTAGGG |
| Mouse *Ulk1* sg3 Rev | AAACCCCTAGTGGTACGCAGACTAC |

Fwd, forward primer; Rev, reverse primer

**Table S5. TIDE analysis primers.**

| Name | Sequence 5'→3' |
|---|---|
| RB1CC1 sg1 Fwd | TGGTGTTGCTTTGTAATGCTTC |
| RB1CC1 sg1 Rev | GAATTCAACTTGCATACCTCCC |
| RB1CC1 sg2 Fwd | ATGGAGAGGTGGTGAGATTTGT |
| RB1CC1 sg2 Rev | AGAGAGCACCAGTTCAGTGGAT |
| ATG13 sg1 Fwd | AGACTGTCCAAGTGATTGTCCA |
| ATG13 sg1 Rev | CACAGCAGAAAGTTAAGAACCAAA |
| ATG13 sg2 Fwd | GCTAGAATGTGAAGTTCCCCTC |
| ATG13 sg2 Rev | TAAATAAGCACGTGTGTCAGGG |
| NRBF2 sg1 Fwd | AGCGTGAAGAAAGATTGAAAGC |
| NRBF2 sg1 Rev | TGTTTTATCATCTTTTGGGGCT |
| NRBF2 sg2 Fwd | GTGTTGGGTTATTCACAAAGGA |
| NRBF2 sg2 Rev | GCTTTCAATCTTTCTTCACGCT |
| PIK3C3 sg1 Fwd | GGAATGAATGGCTGAAACTACC |
| PIK3C3 sg1 Rev | GAAAAGGGTCAGAAAGCTGCTA |
| PIK3C3 sg2 Fwd | TGCTCTGTAATCTAGGACGGTG |
| PIK3C3 sg2 Rev | AAATTCCATCAGAAACAGTGCC |
| ATG14 sg1 Fwd | ATAATCGCAAACTTGGTGACCT |
| ATG14 sg1 Rev | CAGGAAAACCAATGACCCTAGA |
| ATG14 sg2 Fwd | TTATGTCTGGTCATGGAGCATC |
| ATG14 sg2 Rev | TTCAATTACTTGCCTGTTGCAG |
| NDP52 sg1 Fwd | ACCTTCATGTGGGTTACTTTGC |
| NDP52 sg1 Rev | CATCCTCATCCACATAGCAGAA |
| NDP52 sg2 Fwd | GGATTTTGCTTTTCCTGACTTG |
| NDP52 sg2 Rev | ATGAAATGCTGGGTGAAGGTAT |
| SQSTM1 sg1 Fwd | GAAGGTGAAACACGGACACTTC |
| SQSTM1 sg1 Rev | GGTATCCTGAATTCTTGCCTTG |
| SQSTM1 sg2 Fwd | CTACGACTTGTGTAGCGTCTGC |
| SQSTM1 sg2 Rev | AGTTTCCTGGTGGACCCATT |
| NBR1 sg1 Fwd | GGAGCAGGCTAGAGACTTTGTT |
| NBR1 sg1 Rev | AGTTAAAACCCAAGCGAGACAG |
| NBR1 sg2 Fwd | TGTATCTGTGGAGTTCATTGCC |
| NBR1 sg2 Rev | TGTCAGTCAATGCTCACCTCTT |
| TAP1 sg1 Fwd | CTCATCACTTGGAACCTGTCTG |
| TAP1 sg1 Rev | GGTACCATTTTCCCACCTTCTT |
| TAP1 sg2 Fwd | AGTACTGCTACTTCTCGCCGAC |
| TAP1 sg2 Rev | ATGAGATCAGCTCTCGGAACA |
| TAP2 sg1 Fwd | CATCTCCCTCCCCTCTTATTCT |
| TAP2 sg1 Rev | TTAGTCTCCTGGAAGAAACCGA |
| TAP2 sg2 Fwd | CAAATTGGAACACTGGGGTATT |
| TAP2 sg2 Rev | GTCGGTCCATGTAGGAGAAAAC |
| TAPBP sg1 Fwd | GCAGGTCACCAGACATACAAAC |
| TAPBP sg1 Rev | ACTGAGATAGAGCTCAGGGTCG |
| TAPBP sg2 Fwd | TCCTTCTCTACACTCAGACCCC |
| TAPBP sg2 Rev | ATATGCTGACCATCAGCCAAG |
| Mouse *Ulk1* sg2 Fwd | GGGGTAGTAATGACACCACCTC |
| Mouse *Ulk1* sg2 Rev | ACTTCTCGAATCTCCCAAACAA |

Fwd, forward primer; Rev, reverse primer

**Table S6. TIDE analysis results**

| Cell Line | sgRNA | TIDE efficiency |
|---|---|---|
| SCC90 | ATG13 sg1 | 76.8 |
| | ATG13 sg2 | 81.7 |
| | ATG14 sg1 | 90.6 |
| | ATG14 sg2 | 98.0 |
| | TAPBP sg1 | 86.9 |
| | TAPBP sg2 | 96.0 |
| SCC152 | ATG13 sg1 | 75.1 |
| | ATG13 sg2 | 86.8 |
| | ATG14 sg1 | 92.3 |
| | ATG14 sg2 | 56.8 |
| | RB1CC1 sg1 | 44.1 |
| | RB1CC1 sg2 | 82.1 |
| | PIK3C3 sg1 | 90.4 |
| | PIK3C3 sg2 | 85.5 |
| | NRBF2 sg2 | 75.1 |
| | NDP52 sg1 | 83.6 |
| | NDP52 sg2 | 77.4 |
| | SQSTM1 sg1 | 87.3 |
| | SQSTM1 sg2 | 98.3 |
| | NBR1 sg1 | 76.2 |
| | NBR1 sg2 | 78.5 |
| | TAPBP sg1 | 80.1 |
| | TAPBP sg2 | 93.5 |

TIDE, Tracking of Indels by DEcomposition

**Table S7. Antibodies**

| Antibody | Reactivity | Source | Catalog | Clone No. | RRID |
|---|---|---|---|---|---|
| HLA Class I ABC | Human | ProteinTech | 15240-1-AP | | AB_1557426 |
| HLA Class I ABC | Human | ProteinTech | 66013-1-1g | 5C5B7 | AB_11042593 |
| Cytokeratin (pan-reactive) | Human | BioLegend | 628604 | C-11 | AB_2563652 |
| LC3B I/II | Human, Mouse | Cell Signaling | 2775S | | AB_915950 |
| ATG14/Barkor | Human, Mouse | ProteinTech | 19491-1-AP | | AB_10642701 |
| Phospho-ATG14 (Ser29) | Human, Mouse | Cell Signaling | 92340 | D4B8M | AB_2800182 |
| β-actin | Human, Mouse | BioLegend | 664804 | W16197A | AB_ 2728496 |
| ULK1 | Human, Mouse | Cell Signaling | 8054 | D8H5 | AB_11178668 |
| NBR1 | Human, Mouse | ProteinTech | 16004-1-AP | | AB_2251178 |
| P62 | Human | ProteinTech | 18420-1-AP | | AB_10694431 |
| NDP52 | Human | ProteinTech | 12229-1-AP | | AB_11182600 |
| Calreticulin | Human, Mouse | ProteinTech | 27298-1-AP | | AB_2880835 |
| HLA-A, B, C FITC | Human | BioLegend | 311404 | W6/32 | AB_314873 |
| CD11b eFLuor 450 | Mouse | eBioScience | 48-0112-80 | M1/70 | AB_1582236 |
| Ly6C BV510 | Mouse | BioLegend | 128033 | HK1.4 | AB_2562351 |
| F4/80 BV605 | Mouse | BioLegend | 123133 | BM8 | AB_2562305 |
| CD3 BV711 | Mouse | BioLegend | 100241 | 17A2 | AB_2563945 |
| CD45 PerCP | Mouse | BioLegend | 103129 | 30-F11 | AB_893343 |
| I-A/I-E PE | Mouse | BioLegend | 107608 | M5/114.15.2 | AB_313323 |
| NKp46 PE/Cy7 | Mouse | BioLegend | 137617 | 29A1.4 | AB_11219186 |
| CD19 APC | Mouse | BioLegend | 115512 | 6D5 | AB_313647 |
| Ly6G AF700 | Mouse | BioLegend | 127621 | 1A8 | AB_10643269 |

| | | | | | |
|---|---|---|---|---|---|
| CD4 FITC | Mouse | BioLegend | 100510 | RM4-5 | AB_312713 |
| CD8a PE/Cy7 | Mouse | BioLegend | 100722 | 53-6.7 | AB_312761 |
| CD62L AF700 | Mouse | BioLegend | 104426 | MEL-14 | AB_493719 |
| CD44 PE | Human, Mouse | BioLegend | 103007 | IM7 | AB_312958 |
| CD28 | Mouse | BioLegend | 102102 | 37.51 | AB_312867 |

RRID, Research Resource Identifier
